# Multiscale model of primary motor cortex circuits predicts in vivo cell type-specific, behavioral state-dependent dynamics

**DOI:** 10.1101/2022.02.03.479040

**Authors:** Salvador Dura-Bernal, Samuel A Neymotin, Benjamin A Suter, Joshua Dacre, Julia Schiemann, Ian Duguid, Gordon MG Shepherd, William W Lytton

## Abstract

Understanding cortical function requires studying multiple scales: molecular, cellular, circuit and behavior. We developed a biophysically detailed multiscale model of mouse primary motor cortex (M1) with over 10,000 neurons and 30 million synapses. Neuron types, densities, spatial distributions, morphologies, biophysics, connectivity and dendritic synapse locations were tightly constrained by experimental data. The model includes long-range inputs from 7 thalamic and cortical regions, as well as noradrenergic inputs from locus coeruleus. Connectivity depended on cell class and cortical depth at sublaminar resolution. The model accurately predicted in vivo layer- and cell type-specific responses (firing rates and LFP) associated with behavioral states (quiet wakefulness and movement) and experimental manipulations (noradrenaline receptor blocking and thalamus inactivation). It also enabled evaluation of multiple mechanistic hypotheses underlying the observed activity. This quantitative theoretical framework can be used to integrate and interpret M1 experimental data and sheds light on the cell type-specific multiscale dynamics associated with a range of experimental conditions and behaviors.

## Introduction

Understanding cortical function requires studying its components and interactions at different scales: molecular, cellular, circuit, system and behavior. Biophysically detailed modeling provides a tool to integrate, organize and interpret experimental data at multiple scales and translate isolated knowledge into an understanding of brain function. Previous approaches have emphasized structural aspects based on layers and the broad classification of excitatory and inhibitory neurons (***Potjans and Diesmann, 2014; Douglas et al., 1989***). Modern anatomical, physiological and genetic techniques allow an unprecedented level of detail to be brought to the analysis and understanding of cortical microcircuits (***Luo et al., 2018**; **Adesnik and Naka, 2018***). In particular, several neuron classes can now be identified based on distinct gene expression, morphology, physiology and connectivity. Cortical excitatory neurons are broadly classified by their axonal projection patterns into intratelencephalic (IT), pyramidal-tract (PT) and corticothalamic (CT) types (***Greig et al., 2013; Harris and Shepherd, 2015; Zeng and Sanes, 2017***). Recent research has also revealed that connections are cell-type and location specific, often with connectivity differences at different cortical depths within layers (***Anderson et al., 2010; Brown and Hestrin, 2009; Morishima and Kawaguchi, 2006***).

Primary motor cortex (M1) plays a central role in motorcontrol, butto date M1 circuits have only been modeled to a limited extent (***Chadderdon et al., 2014; Neymotin et al., 2016b; Heinzle et al., 2007; Morita and Kawaguchi, 2015; Hoshino et al., 2019***). We and others have extensively studied mouse M1 circuits experimentally, and characterized cell subclasses and many cell-type and sublaminar-specific local and long-range connections (***Papale and Hooks, 2017; Shepherd, 2009; Kaneko,2013; Morishima et al., 2011***). A major focus of these anatomical and physiological studies has been the distinct cell classes of layer 5 (L5): L5B PT cells – the source of the corticospinal tract, and other pyramidal tract projections, and L5 IT cells which project bilaterally to cortex and striatum. Morphology and physiology differ across the two types. L5 IT cells are thin-tufted and show spike frequency adaptation. L5B PT cells are thick-tufted and show little spike frequency adaptation, but strong sag potentials. Their spiking dynamics in vivo have also been shown to differ (***Saiki et al., 2018***). In terms of their synaptic interconnectivity these types exhibit a strong asymmetry: connections go from IT to PT cells, but not in the opposite direction (***Kiritani et al., 2012; Morishima and Kawaguchi, 2006***). The strength of their local excitatory input connections is also dependent on PT position within layer 5B, with cells in the upper sublayer receiving the strongest input from layer 2/3 (***Anderson et al., 2010; Hooks et al., 2013; Yu et al., 2008; Weiler et al., 2008***). These and several other highly specific local and long-range wiring patterns are likely to have profound consequences in terms of understanding cortical dynamics, information processing, function and behavior (***Li et al., 2015b***).

A key unanswered question in the motor system, and more generally in neural systems (***Mott et al., 2018; Hsu et al., 2020; Getting, 1989***), is how cell and circuit dynamics relate to behavior. Both IT and PT cell types play a role in motor planning and execution and both have been implicated in motor-related diseases (***Shepherd, 2013***). We have previously shown that the hyperpolarization-activated current (*I*_h_), a target of noradrenergic neuromodulation, is highly expressed in PT cells and affects its synaptic integration and electrophysiological properties (***Sheets et al., 2011; BICCN, 2021***). In vivo studies also reveal noradrenergic neuromodulatory inputs from locus coeruleus (LC) and long-range inputs from thalamus and cortex causally influence M1 activity and behavioral states (***Boychuk et al., 2017; Schiemann et al., 2015; Guo et al., 2021***). Specifically, blocking noradrenergic input to M1 impaired motor coordination (***Schiemann et al., 2015***), and disrupting the cerebellar-recipient motor thalamus projections to M1 can impair dexterity (***Guo et al., 2021***) or block movement initiation (***Dacre et al., 2021**).* These modulatory and long-range projections have been shown to be cell type-specific, and characterized in ex vivo slice experiments (***Sheets et al., 2011; Yamawaki and Shepherd, 2015; Suter and Shepherd, 2015***), but how these relate to in vivo activity, including the exact cellular and circuit mechanisms underpinning behavioral statedependent M1 activity, remains largely unknown. A biologically realistic model of M1 can be used to address this current knowledge gap by generating hypotheses and predictions relating circuit dynamics to function and behavior.

Previous models of M1 circuits are scarce and lack the detail across scales requied to adequately address these questions. The M1 models by ***Morita and Kawaguchi (2015); Hoshino et al. (2019)*** only included a single layer with two cell types. ***Heinzle et al. (2007)*** proposed a microcircuit model of the frontal eye field with 4 layers and multiple cell types. However, all of these circuit models included highly simplified neuron models with limited biophysical detail and no morphological detail. Our previous work modeling M1 (***Chadderdon et al., 2014; Neymotin et al., 2016b**)* incorporated neuron models with 5-compartment morphologies and multiple ionic channels, as well as several cell types distributed across 5 cortical layers and connected based on layer and cell type. However, it lacked neuron models tuned to cell type-specific electrophysiological data, realistic neuronal densities, noradrenergic and long-range inputs, and certain connectivity details, including depth-dependence and subcellular distribution of synapses.

We have now developed a multiscale model of mouse M1 incorporating recent experimental data and reproducing in vivo layer- and cell type-specific behavior-dependent responses. The model simulates a cylindric cortical volume with over 10 thousand neurons and 30 million synapses. We attempted, as far as possible, to base parameters on data obtained from a single species, strain and age range, and from our own experimental work. However, these data are necessarily incomplete, and we have therefore combined additional data from multiple other sources. We focused particularly on the role of L5 excitatory neurons, utilizing detailed models of layer 5 IT and PT neurons with full dendritic morphologies of 700+ compartments based on anatomical cell reconstruction and ion channel distributions optimized to in vitro experimental measures. The task of integrating experimental data into the model required us to develop several novel methodological techniques for network simulation design, including: 1) specifying connections as a function of normalized cortical depth (NCD) – from pia to white matter – instead of by layer designations, with a 100-150 *μm* resolution; 2) identifying and including specific dendritic distributions associated with particular inputs using features extracted from subcellular Channelrhodopsin-2-Assisted Circuit Mapping (sCRACM) studies (***Hooks et al., 2013; Suter and Shepherd, 2015***); and 3) utilizing a high-level declarative modeling tool, NetPyNE, to develop, simulate, optimize, analyze and visualize the model (***Dura-Bernal et al., 2019***).

Our M1 model exhibited neuronal firing rates and oscillations that depended on cell class, layer and sublaminar location, and behavioral state, consistent with in vivo M1 data. Behavioral changes (quiet wakefulness vs movement) were modeled by modifying noradrenergic inputs from LC and motor thalamus inputs. Our cortical model also captured the effects of experimental manipulations, including blocking of norardrenergic receptors and motor thalamus inactivation. The model provided different multiscale mechanistic hypotheses for the observed behavioral deficits, linking noradrenaline blockade to cell type specific changes in *I*_h_ and/or potassium conductances and the subsequent changes in neuronal firing patterns. The simulations generated experimentally-testable quantitative predictions about layer- and cell type-specific responses for the different behavioral states and experimental manipulations. Two key model predictions were that stronger thalamic and noradrenergic inputs are required to activate the deeper (associated with motor execution) vs superficial L5B PT neurons, and that L5 interneurons support switching between PT and IT output through mutual disynaptic inhibition. Simulations also shed new light on M1 circuitry and biophysical mechanisms associated with dynamic aspects of behavior-related activity, including PT cells predominantly mediating an increase in gamma physiological oscillations recorded in L5 local field potentials during movement. We are making our model freely available as a community resource so that others can update and extend it, incorporating new data such as that from the M1 multimodal cell census and atlas recently released by the BRAIN Initiative Cell Census Network (***BICCN, 2021***).

## Results

### Overview of model development and simulations

We implemented a biophysically-realistic model of the mouse M1 microcircuit representing a cylindrical volume of 300 *μm* diameter (Fig. 1). The model included over 10,000 neurons with 35 million synapses. Cell properties, locations, and local and long-range connectivity were largely derived from a coherent set of experimental data. Available experimental data was particularly detailed for two L5 populations that were the focus of this study: pyramidal tract (PT) corticospinal cells and intratelencephalic (IT) corticostriatal cells. One innovative feature in the network presented here was the inclusion of a layer 4 for motor cortex, consistent with its recent characterization ***(Yamawaki et al., 2015; Bopp et al., 2017; Barbas and García-Cabezas, 2015; BICCN, 2021***). The model was developed using the NetPyNE (***Dura-Bernal et al., 2019***) modeling tool and the NEURON simulation engine (***Carnevale and Hines, 2006***). Over 20,000 simulations were required to progressively construct and improve the model. Simulations required over 8 million high performance computing (HPC) cluster core-hours to arrive at the results shown, primarily during model building. One second of simulation (model) time required approximately 96 core-hours of HPC time. We employed a grid search on underconstrained connectivity parameters – e.g. inhibitory to excitatory weight ratios – to identify simulations that produced physiologically realistic firing patterns across populations.

**Figure 1.**
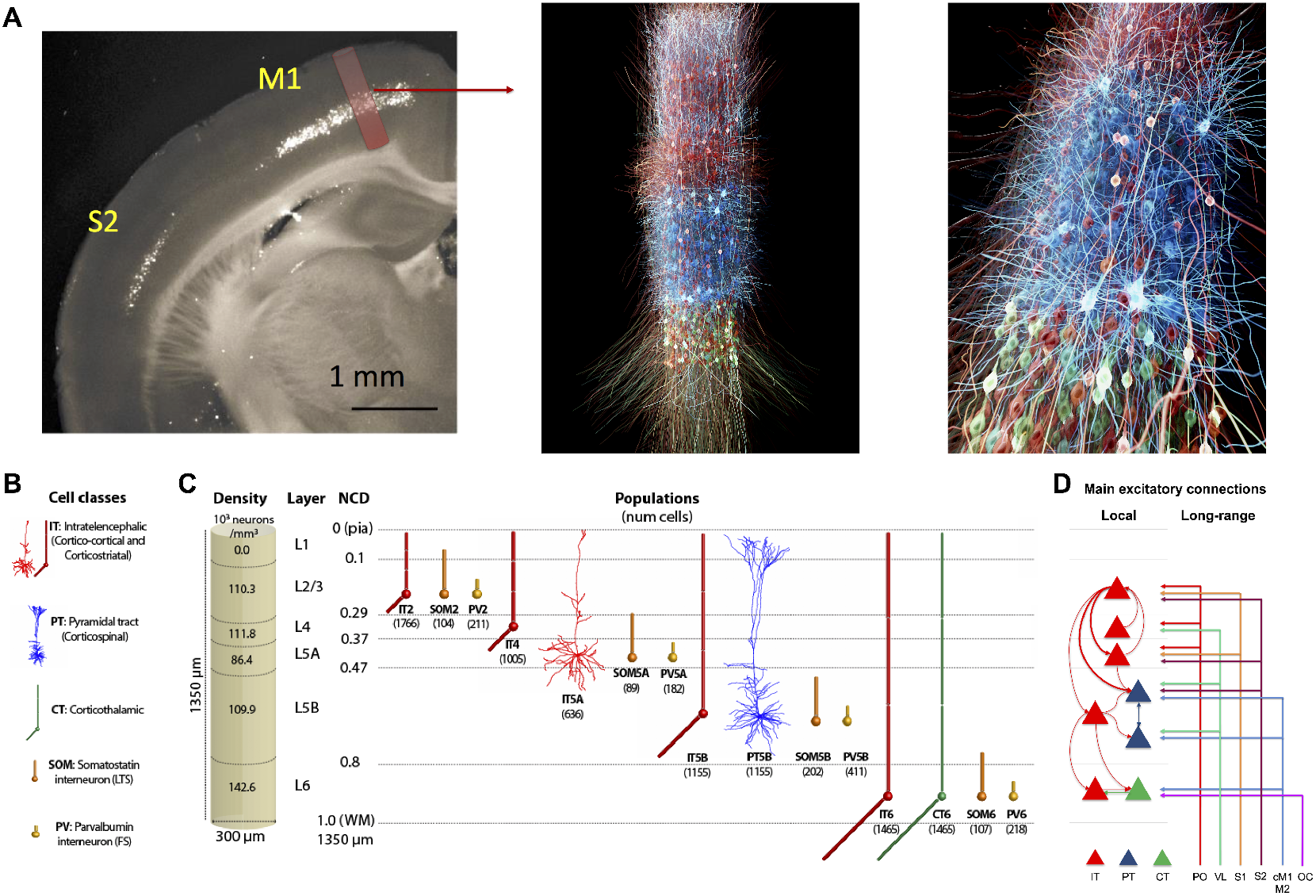
M1 microcircuit model: 3D visualization, connectivity, dimensions, neuronal densities, classes and morphologies. **A.** *left panel:* Epifluorescence image of coronal brain slice of mouse showing M1 and S1 regions, with approximate anatomical location and area of simulated cylindrical tissue (adapted from (***Suter et al., 2013**)). middle and right panels>:* 3D visualization of M1 network, showing location and stylized morphologies of 20% of excitatory IT (red), PT (blue) and CT (green) cells, and snapshot of simulated activity with spiking neurons in brighter color (visualization by nicolasantille.com). **B.** Cell classes modeled. IT5A and PT5B neurons are simulated in full morphological reconstructions. Other excitatory types and inhibitory neurons use simplified models with 2-6 compartments. All models are conductance-based with multiple ionic channels tuned to reproduce the cell’s electrophysiology. **C.** Dimensions of simulated M1 cylindrical volume with overall cell density per layer designation (left), and cell types and populations simulated (right). **D.** Schematic of main local and long-range excitatory connections (thin line: medium;thick line: strong). Note the unidirectional projections from ITs to PTs, with a particularly strong projection arising from L2/3. (IT: intratelencephalic cells – corticostriatal;PT: pyramidal-tract cells – corticospinal;CT corticothalamic cells. PO: posterior nucleus of thalamus;VL: ventrolateral thalamus;S1: primary somatosensory;S2: secondary somatosensory;cM1: contralateral M1;M2: secondary motor;OC: orbital cortex;PV: parvalbumin basket cells, SOM: somatostatin interneurons;number of cells in each population shown in brackets;left shows L1–L6 boundaries with normalized cortical depth – NCD from 0 = pia to 1 = white matter.)

As expected from results in other systems, there was no single “right” model that produced these realistic firing patterns but rather a family of models (degenerate parameterization) that were within the parameter ranges identified by experiment (***Golowasch et al., 2002; Prinz and Marder, 2003; Edelman and Gally, 2001; Ratté and Prescott, 2016***). From these, we selected one *base model,* representing a single parameter set, to illustrate in this paper. This base model was tested for robustness by changing randomization settings to provide a *model set,* with analysis of raw and average data from 25 simulations: 5 random synaptic input seeds × 5 random connectivity seeds (based on connectivity density). This can be considered analogous to testing multiple trials and subjects in an experimental setup. The full model set showed qualitatively similar results with low variance in bulk measures (population rates, oscillation frequencies) for changes in randomization settings.

We used the base model and model set to characterize firing and local field potential (LFP) patterns in response to different levels of long-range inputs and noradrenergic (NA) neuromodulation associated with different behavioral states and experimental manipulations of mouse M1 in vivo (***Schiemann et al., 2015***) (see Table 1). Long-range inputs originated from seven regions: posterior nucleus of thalamus (PO), ventrolateral thalamus (VL), primary somatosensory cortex (S1), secondary somatosensory cortex (S2), contralateral M1 (cM1); secondary motor cortex (M2),and orbital cortex (OC). In the context of this model, VL will be equivalent to the motor thalamus (MTh), for consistency with the experimental study (***Schiemann et al., 2015***). The two behavioral states corresponded to *quiet* wakefulness and self-paced, voluntary *movement.* Each of these states was simulated under three different experimental manipulations mimicking those previously performed in vivo (***Schiemann et al., 2015***): *control,* motor thalamus inactivation (*MTh inactivation)* and blocking input from locus coeruleus (LC) via noradrenergic receptor antagonists (*NA-R block).* The effect of changes in noradrenergic neuromodulation, driven by inputs from LC, were simulated by altering *I*_h_ conductance in PT cells (see Table 1 and Methods), consistent with in vitro findings ***(Sheets et al., 2011)**.* Results are presented both in terms of cell class and cell population. We focused on three excitatory classes: intratelencephalic (IT), pyramidal-tract (PT), corticothalamic (CT); and two inhibitory classes: parvalbumin-expressing fast-spiking basket cells (PV), somatostatin-expressing low-threshold spiking cells (SOM). Cell populations are defined by both class and layer (e.g. IT5A indicates class IT in layer 5A; CT6 is class CT in layer 6). We use our results to explain and predict the response of M1 circuitry under the different behavioral states and experimental manipulations simulated.

**Table 1.** Motor thalamus (MTh) input and noradrenergic (NA) input associated with the different experimental manipulations and behavioral states simulated in the M1 model. NA input is modeled by modifying the conductance of PT *T*_h_.

| Experimental manipulation | Behavioral State | MTh input | NA input (PT $I_h$) |
| --- | --- | --- | --- |
| Control | Quiet | Low (0-2.5 Hz) | Low NA (75% $I_h$) |
| Control | Movement | High (0-10 Hz) | High NA (25% $I_h$) |
| MTh inactivation | Quiet | Very low (0-0.01 Hz) | Low NA (75 % $I_h$) |
| MTh inactivation | Movement | Very low (0-0.1 Hz) | High NA (25% $I_h$) |
| NA-R block | Quiet | Low (0-2.5 Hz) | Very low (100% $I_h$) |
| NA-R block | Movement | High (0-10 Hz) | Very low (100% $I_h$) |

We characterized in vivo spontaneous activity in the base model. This was simulated based on expected background drive of ≤5 Hz from all long-range inputs, and low NA input resulting in medium level *I*_h_ (75%) in PT cells (Fig. 2) (***Yamashita et al., 2013; Hirata and Castro-Alamancos, 2006***). These properties were consistent with the quiet wakefulness state and control conditions as recorded by whole-cell patch-clamp electrophysiology in awake mice in vivo ***(Schiemann et al., 2015)**.* We validated the M1 model cell type- and layer-specific firing rates against available in vivo experimental data from mouse motor cortex (***Schiemann et al., 2015; Zagha et al., 2015; Li et al., 2016; Estebanez et al., 2018; Economo et al., 2018***) (Fig. 2*B*). All population mean and median firing rates ranged between 0.1 and 10 Hz, and maximum rates (excluding outliers) were below 35 Hz, for both model and experiment. More specifically, we compared L2/3 IT (median±IQR model=1.8 ± 4.0 Hz, exp=0.3 ± 0.7 Hz), L5B IT (model=6.5 ± 8.8 Hz, exp=3.2 ± 2.5 Hz), L5B PT (model=1.8 ± 4.8 Hz, exp=4.6 ± 4.6 Hz). Since certain studies did not distinguish between cell types or sublayers we also compared L5B IT/PT(model=4.8±8.5 Hz, exp=5.1±6.0 Hz) and L5 IT/PT(model=5.5±9.2 Hz, exp1=1.7± 4.0 Hz, exp2=7.6 ± 8.5 Hz, exp3=2.4 ± 4.7 Hz). Significant statistical differences among population firing rates from different studies are expected, and therefore these were also expected between model and experiment. An example is L5 IT/PT where two experimental datasets were statistically significantly different (exp1=1.7±4.0 Hz, exp2=7.6±8.5 Hz;p = 6.2e-15, rank-sum test), whereas this was not the case when comparing the L5 IT/PT model to experiment (model=5.5±9.2 Hz, exp2=7.6± 8.5 Hz *p* = 0.43, rank-sum test). Overall, these results indicate that the range of firing rates and variability in the model was consistent with that of in vivo mouse data.

**Figure 2.**
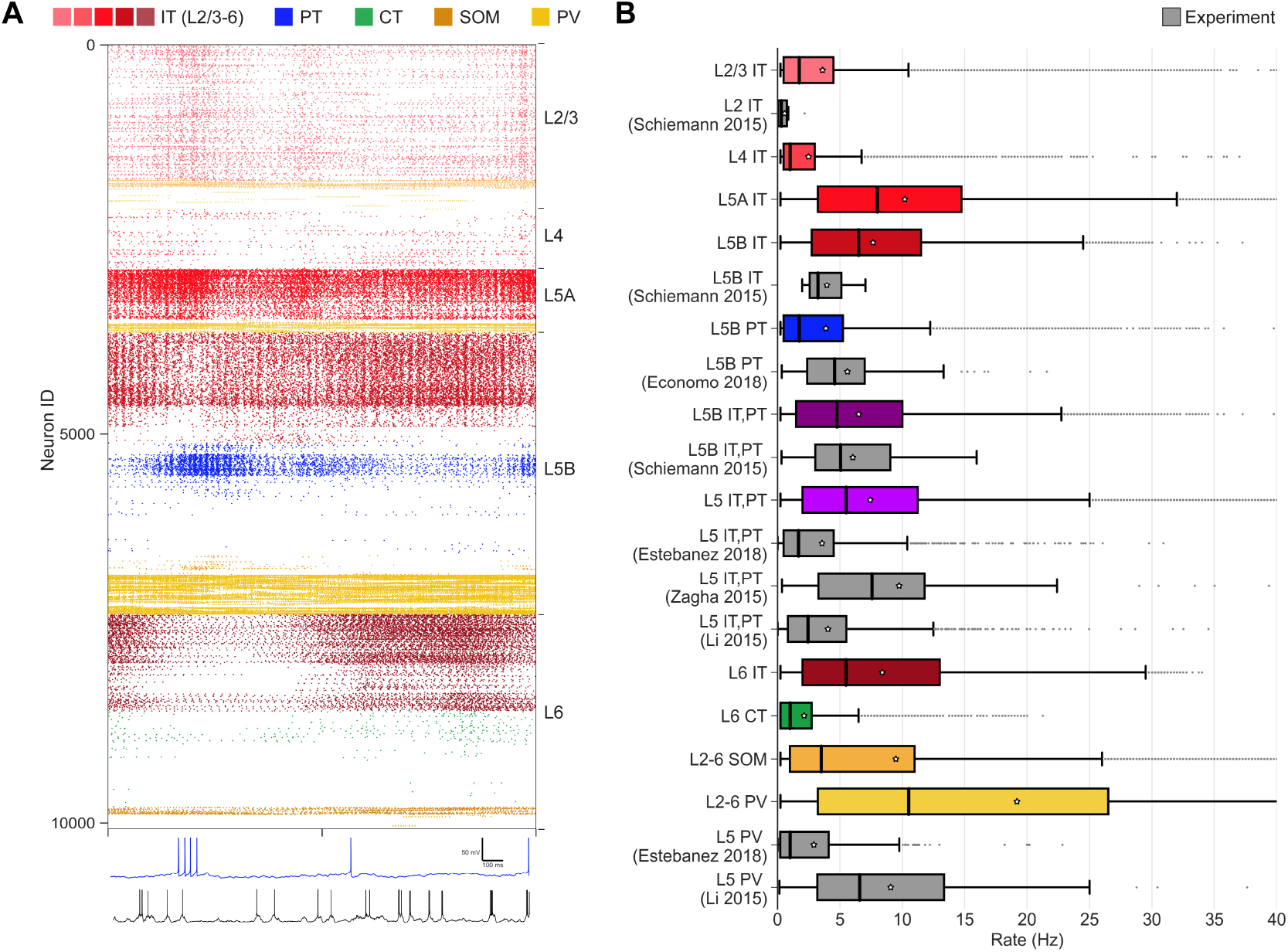
M1 cell type and layer-specific firing dynamics during quiet wakefulness state and control condition (spontaneous activity) The quiet state was simulated by driving the network with background activity (≤ 5 Hz) from all long-range inputs, and medium level *I*_h_ (75 %) in PT cells (low NA modulation). **A.** *Top:* Raster plot of mid-simulation activity (2 s of base model simulation shown;cells grouped by population and ordered by cortical depth within each population). *Bottom:* Example model (blue) and experiment (black) PT5B voltage traces. **B.s** Firing rates statistics (boxplots) for different cell types and layers in the model set (color bars) and experiment (gray bars).

Activity patterns were not only dependent on cell class and cortical-layer location, but also sub-laminar location. This supports the importance of identifying connectivity and analyzing activity by normalized cortical depth (NCD) in addition to layer. For example, L5B PT firing rates decreased with cortical depth (Fig. 2*A*), consistent with depth-weighted targeting from L2/3 IT projections ***(Anderson et al., 2010; Weiler et al., 2008**).* This pattern of firing was consistent across network varia-tionswith different wiring and input randomization seeds. L5A/B IT exhibited similar cortical-depth dependent activity. L2/3 and L4 IT populations showed overall lower rates than L5 IT, consistent with weaker excitatory projections onto these populations from local M1 (***Weiler et al., 2008; Yamawaki et al., 2015)**,* and from long-range inputs (***Mao et al., 2011; Suter and Shepherd, 2015; Yamawaki et al., 2015)**.* In particular, the main source of L4 IT input was thalamic, in correspondence with the well-described pattern in sensory cortex (***Yamawaki et al., 2015**).* Despite the weaker response, L2/3 IT showed slow oscillatory activity around delta frequency. Within L6, superficial cells of IT and CT populations were more active than deeper ones. This was due to stronger intralaminar, L5B IT (***Weiler et al., 2008; Yamawaki and Shepherd, 2015**)* and long-range inputs, primarily from orbital and contralateral motor cortices (for more details on model connectivity see Methods Fig. 8) (***Hooks et al., 2013**).* Weaker local projections onto L6 CT compared to L6 IT resulted in firing rate differences between CT and IT. Although the model anatomical connectivity was empirically constrained, population responses are not fully defined by the anatomy, but emerge from the complex dynamical interplay across different excitatory and inhibitory populations.

### M1 firing dynamics during movement

The model reproduced experimental cell type-specific dynamics associated with movement. The movement state was simulated by increasing long-range inputs from ventrolateral thalamus (VL; here equivalent to motor thalamus, MTh) to 0-10 Hz (uniform distribution), and reducing *I*_h_ conductance to 25% in PT cells, to simulate high NA neuromodulatory inputs from LC. The remaining 6 long-range inputs (PO, S1, S2, cM1, M2, OC) continued to provide background drive (≤ 5Hz). This resulted in a large increase in L5B PT activity and the development of a strong gamma oscillation (observable in the spiking raster activity Fig. 3A). PT5B_lower_ neurons, which were largely silent during the quiet state, now exhibited similar activity to PT5B_upper_. This is consistent with the involvement of PT, and particularly PT5B_lower_ (***Economo et al., 2018**),* in motor control. During movement, the activity of L2/3 IT and L5 IT decreased moderately, whereas L4 IT, L6 IT and L6 CT firing rates remained similar. There was a transition period from quiet to movement that lasted approximately 500ms, during which there was a peak in the activity of L5 IT and PT5B_upper_, consistent with efferent motor thalamic projections. This transitory activity peaks could also be seen in most of the remaining model set simulations. Although IT2/3 exhibited a similar transition peak in the base model, this was not apparent in other model set simulations, suggesting this could have resulted from the ongoing L2/3 IT delta oscillations.

**Figure 3.**
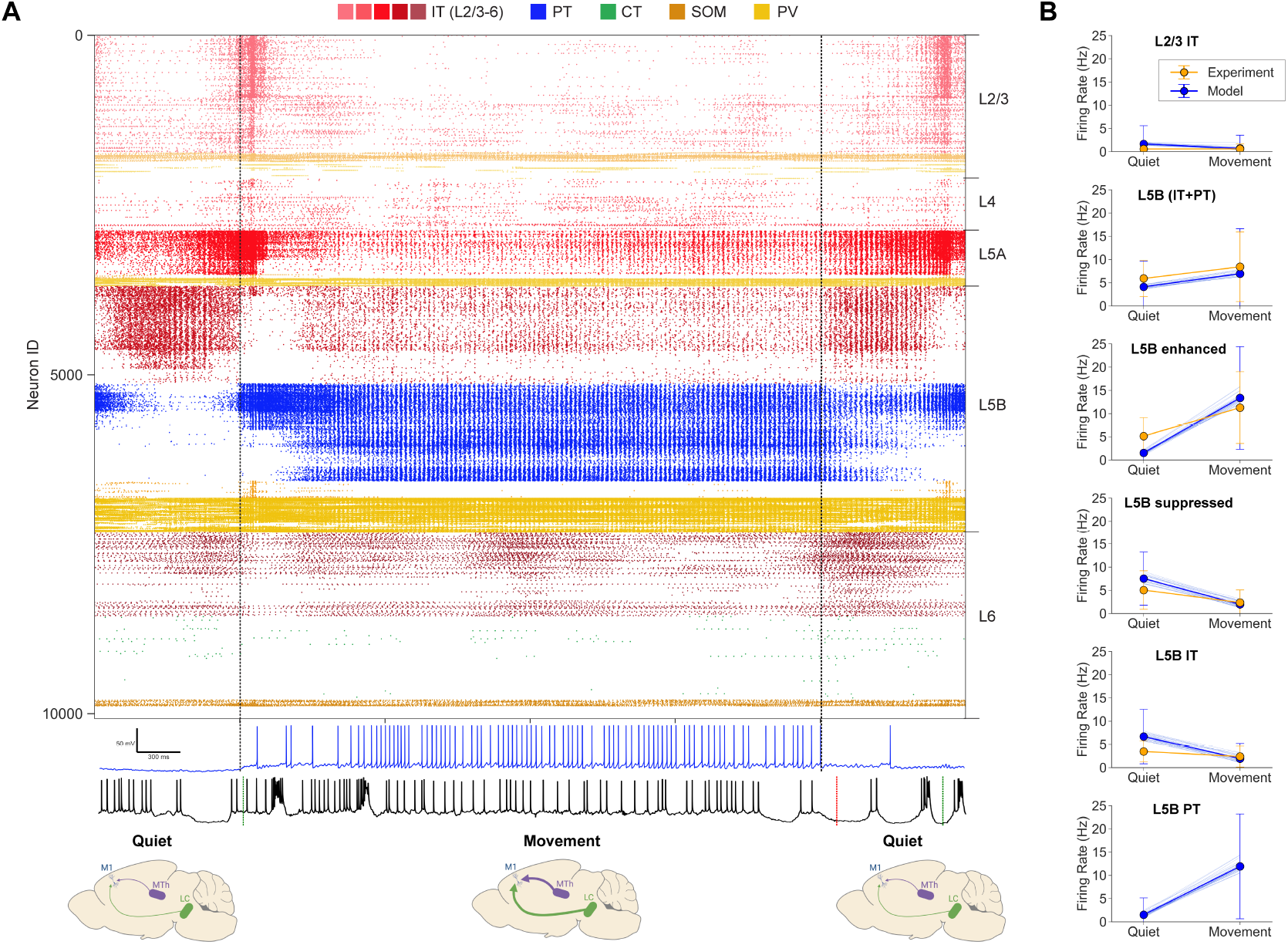
M1 cell-type and layer-specific firing dynamics during quiet and movement states under the control condition. The movement state was simulated by driving the network with increased activity (0-10Hz) from motor thalamus, background activity (≤5Hz) from the 6 remaining long-range inputs, and reducing *I*_h_ to 25% in PT cells (mimicking high NA modulation). **A.** *Top:* Raster plot of activity transitioning from quiet (1s) to movement (4s) to quiet (1s) states (6s of base model simulation shown;cells grouped by population and ordered by cortical depth within each population). *Bottom:* Example model PT5B (blue) and experiment (black) voltage traces. **B.** Firing rate (mean±SD) in different cell populations for model set (blue) and experiment (orange). Model set includes cell rates of all 25 simulations;the mean rates of each individual simulation shown as thin blue lines. Statistics were computed across 4s for each state.

Model firing rate distributions were generally consistent with experimental data across populations and behavioral states. We compared the quiet and movement population firing rates of the model set against M1 in vivo experimental data (***Schiemann et al., 2015**)* (Fig. 3B). Both model and experiment L2/3 IT cells exhibited low firing rates during both quiet (mean±SD model: 1.6 ± 3.9 Hz; exp: 0.6±0.7 Hz) and movement states (mean±SD model: 0.7±2.8 Hz; exp: 0.6±1.1 Hz). The L5B rates, including both IT and PT, were similar in model and experiment and exhibited a similar increase from quiet (model 4.1 ± 5.5 Hz; exp 5.9 ± 3.9 Hz) to movement (model: 6.9 ± 9.7 Hz; exp: 8.4 ± 7.5 Hz). Following the experimental study data analysis and classification of populations ***(Schiemann et al., 2015)**,* we compared rates of cells that exhibited *enhanced* or *suppressed* activity from quiet to movement. Both L5B_enhanced_ and L5B_suppressed_ rates exhibited comparable trends in model and experiment. The quiet state L5B_enhanced_ mean±SD rates were higher in the model than experiment (model: 1.5±3.6 Hz, exp: 5.1±4.0 Hz) but increased to a similar rate during movement (model: 13.2±11.1 Hz, exp: 11.3 ±7.7 Hz). L5B_suppressed_ model and experiment rates exhibited a similar decrease from quiet (model: 7.5 ±5.7 Hz, exp: 5.0 ±4.2 Hz) to movement states (model: 2.0 ±3.1, exp: 2.3 ±2.7 Hz). L5B IT quiet mean±SD rates were higher for model vs experiment (model: 6.7±5.9 Hz, exp: 3.5±2.3 Hz) but also decreased to a similar level during movement (model: 1.9±3.3 Hz, exp: 2.4±2.3 Hz). Model L5B PT rates increased sharply from quiet (1.5 ±3.6 Hz) to movement (11.9±11.3 Hz). We did not include experiment PT rates in Fig. 3B given their small sample size (N=3) and high variability. However, we note that two of the experiment PT cells showed a decrease from quiet to move (16.0 Hz to 5.6 Hz and 4.7 Hz to 0.6 Hz), and one showed a similar sharp increase to that of the model (3.5 Hz to 13.2 Hz). The robustness of the model was evidenced by the small variability across the mean firing rates of the 25 simulations in the model set, each with different randomization seeds (see thin blue lines in Fig. 3B).

### M1 layer 5 LFP oscillations depend on behavioral state

In vivo studies in mouse vibrissal M1 have shown a decrease of L5 LFP slow oscillations (3-5 Hz) and an increase in gamma oscillations (30-50 Hz) during active whisking (***Zagha et al., 2013**).* Here, we investigated whether similar changes were observed in the L5 LFP of mouse M1 during the self-paced, voluntary movement task (***Schiemann et al,, 2015**),* and if those changes were captured by our simulated M1 LFP (Fig. 4). Importantly, the model was not tuned to reproduce the experiment LFP during either quiet or movement states. Despite this, LFP amplitudes were overall similar in model and experiment (order of 500 *μV*). In both experiment and model, the L5 LFP showed weaker slow oscillations (delta) and stronger fast oscillations (gamma) during movement compared to quiet behavioral states, consistent with the previously reported experiments (***Zagha et al., 2013**).* This is illustrated in the raw LFP signal and spectrogram examples for experiment and model (Figure 4A for quiet and 4B for movement). Model L5 LFP was averaged across the signals recorded from simulated extracellular electrodes at 3 depths within L5: 600*μ*m (L5A), 800*μ*m (upper L5B) and 1000*p*m (lower L5B). The experimental LFP dataset was recorded in vivo from L5 extracellular electrodes and preprocessed to remove outliers and potential artifacts (see Methods).

**Figure 4.**
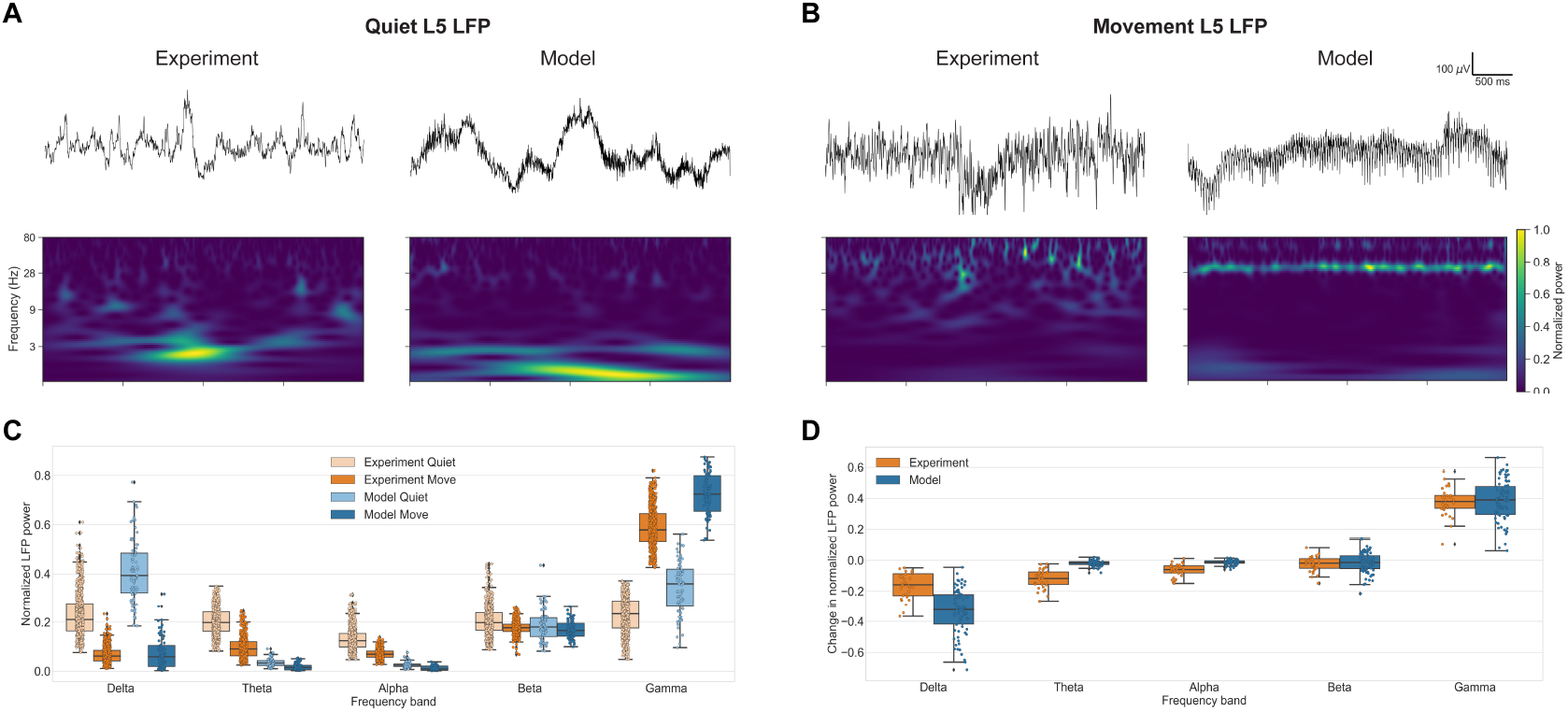
M1 layer 5 LFP oscillations during the quiet and movement states. Example experiment and model raw LFP signals (*top*) and spectrograms (*middle)* during the quiet (**A**) and movement (**B**) states. **C.** Comparison of experiment and model normalized power spectral density (PSD) power across 5 frequency bands during quiet and movement states. **D.** Comparison of experiment and model changes in normalized power spectral density (PSD) power across 5 frequency bands during quiet and movement states.

The model reproduced behavioral-dependent differences across different frequency bands of M1 LFP oscillations. To quantify these differences we calculated the LFP normalized power spectral density (PSD) across the major frequency bands for the experimental and modeling datasets (Fig. 4C). To enable comparison, we segmented the experimental data in 4-second samples, matching the duration of the model dataset samples. Both experiment and model datasets exhibited stronger LFP power at the lower end of the spectrum (delta, theta and alpha bands) during the quiet state, and stronger high-frequency (gamma) LFP power during movement. More specifically, delta (0-4 Hz) power in the quiet state was high in both model vs experiment (median±IQR: model: 0.39±0.16; exp: 0.21±0.11) but decreased to a similar level during movement (model: 0.06±0.09; exp: 0.06 + −0.04). Theta (4-8 Hz) power was overall higher in experiments compared to the model, but in both cases showed higher amplitude in the quiet vs movement states. A similar pattern was observed for the LFP alpha (8-13 Hz) power (model: 0.02±0.01 vs 0.01±0.02; exp:0.12±0.05 vs 0.07±0.03). Beta power (13-30 Hz) remained largely stable from quiet to movement states, and exhibited very similar values for experiment and model (model: 0.18 ± 0.08 and 0.18 ± 0.08; exp: 0.20 ± 0.07 and 0.18 ± 0.03). Gamma power (30-80 Hz) was stronger during movement for both experiment and model (model: 0.36 ± 0.15 and 0.72 ± 0.14; exp: 0.23 ± 0.11 and 0.58 ± 0.11). The increase in PT average firing rates and oscillatory activity depicted in Fig. 3 suggest PT neurons are predominantly responsible for the increase in L5 gamma LFP power.

The model also reproduced the main changes in LFP power from quiet to movement states when looking at paired samples occurring within the same recording. In the previous comparison, the experimental dataset included a larger number of 4-second samples for the quiet (N=3890) than movement (N=2840) states. These were obtained from 30 recordings from different animals, trials and recording sites within L5. In order to more directly quantify the change in LFP power from quiet to movement, we selected the subset of paired 4-second quiet and movement samples that occurred consecutively within the same recording. We then calculated the change in normalized LFP PSD for the resulting 160 pairs of consecutive quiet and movement samples (Fig. 4D). Both model and experiment showed results consistent with the previous analysis: from quiet to movement there was 1) a strong decrease of delta frequency power during movement (model: −0.32±0.19; exp: −0.16±0.14); 2) small changes in theta, alpha and beta power; and 3) large increase in gamma power (model: 0.39 ±0.18; exp: 0.38 ±0.08). These results provide further validation that the model is capturing behavior-related oscillatory dynamics observed in mouse M1 in vivo.

### M1 dynamics during motor thalamus inactivation

To gain insights into the known role of thalamic inputs in regulating M1 output (***Guo et al., 2021; Dacre et al., 2021***) we simulated an experimental manipulation described in our in vivo study ***(Schiemann et al., 2015)**,* consisting of blocking thalamic input to M1 by local infusion of the *GABA_A_* receptor agonist muscimol into the VL/VA complex. Our computational model captured several features of inactivating motor thalamus (MTh) inputs to M1. The MTh inactivation condition was simulated by removing the VL input. The other 6 long-range background inputs (PO, cM1, M2, S1, S2, OC) remained. Under this condition, the change from quiet to movement states only involved reducing and reducing *I*_h_ conductance from 75% to 25% in PT cells, simulating the high NA neuromodulatory inputs from LC. The decrease in movement-associated L5B activity (control: 6.9 ± 9.7 Hz, MTh inact: 4.00±5.7 Hz) after MTh inactivation (Fig. 5A,B) was consistent with that seen experimentally (control: 8.4±7.5 Hz, MTh inact: 2.2±4.0 Hz). The model also captured the strong reduction in the movement-associated L5B_enhanced_ population response following MTh inactivation (model control: 13.3±11.1 Hz, MTh inact: 6.3 ± 7.1 Hz; exp control: 11.3 ± 7.7, MTh inact: 4.2 ± 4.9). The decrease in the model L5B rates was caused by a strong reduction of PT rates (control: 11.9 ± 11.3 Hz, MTh inact: 2.9 ± 6.0 Hz). MTh inactivation resulted in a particularly strong reduction of the movement-associated PT5B_lower_ population, which was practically silenced.

**Figure 5.**
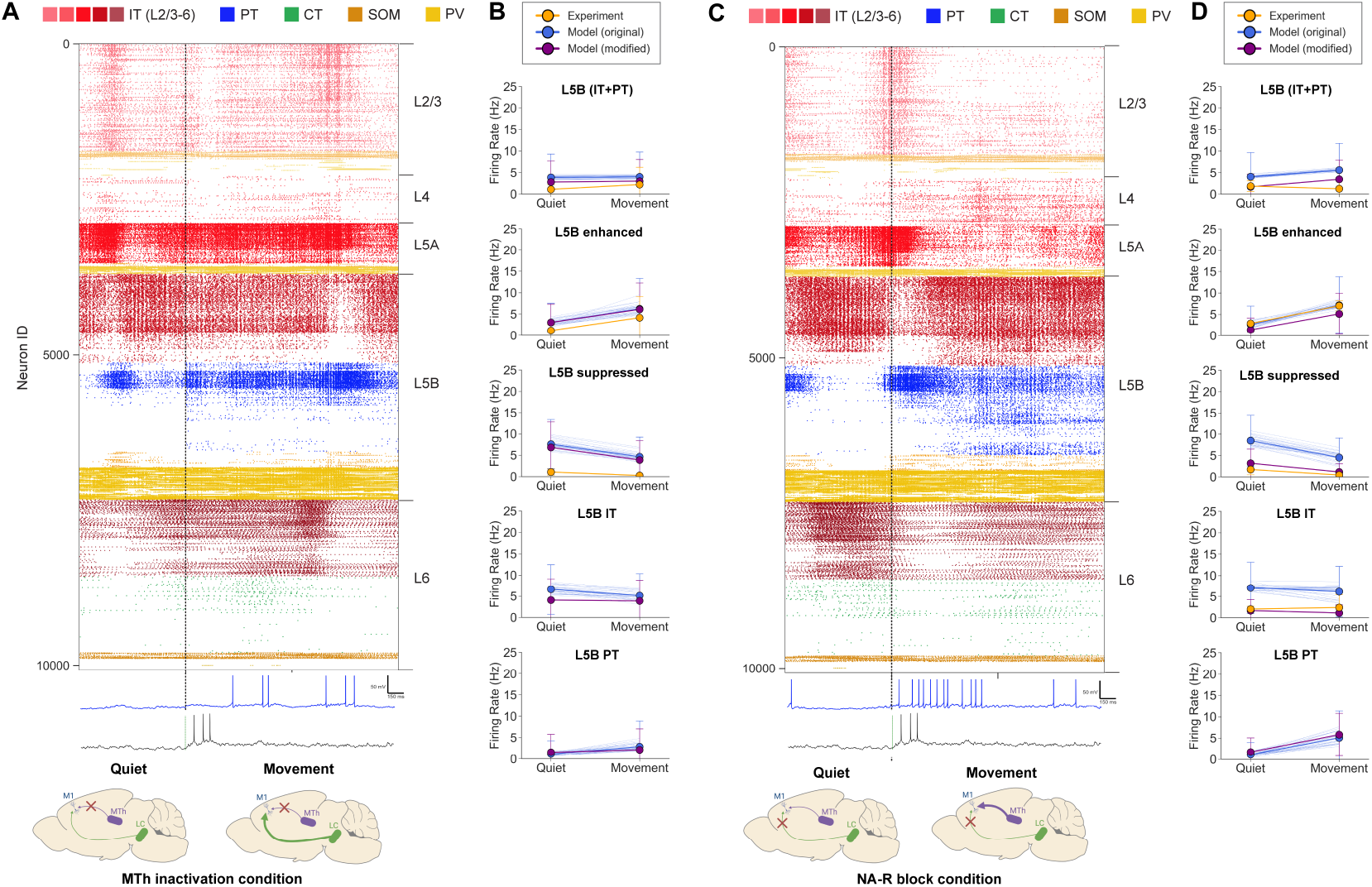
M1 cell-type and layer-specific firing dynamics during the quiet and movement states for the MTh inactivation (A and B) and the NA-R block (C and D) conditions. A. and C. *Top:* Raster plot of activity transitioning from quiet (1s) to movement (2s) (3s of base model simulation shown;cells grouped by population and ordered by cortical depth within each population). *Bottom:* Example model PT5B (blue) and experiment (black) voltage traces. **B. and D.** Firing rate (mean±SD) in different cell populations for the original model set (blue), modified model (purple) and experiment (orange). The modified model decreased long-range inputs from cM1 and M2 for the MTh inactivation condition, and increased K+ conductance for the NA-R block condition. The original model set includes cell rates of all 25 simulations;the mean rates of each individual simulation shown as thin blue lines. Statistics were computed across 4s for each state.

However, results suggested that the model was not adequately capturing some effects of MTh inactivation on M1 L5B, particularly during the quiet state. Specifically, MTh inactivation lead to a reduction of quiet state L5B (control: 5.1 ± 3.9 Hz, MTh inact: 1.1 ± 1.1), as well as L5B_suppressed_, which was not observed in our model, where these two populations rates remained similar. This pointed to future directions to improve our baseline model by evaluating different hypotheses of the mechanisms and circuitry underlying the experimental observations. Here we evaluated one such hypothesis: discrepancies could be due to the lack of interaction between long-range inputs in the model, preventing it from capturing the effects of MTh inactivation on other regions (e.g. M2) that in turn provide input to M1 (see Discussion for more details and alternatives). To evaluate this hypothesis we modified our original model of MTh inactivation by reducing the activity of other cortical long-range inputs (cM1, M2). The modified model better reproduced experimental L5B results (see Fig. 5B purple lines) both for the quiet (orig model: 3.9 ± 5.4; modified model for MTh inactivation: 2.8±4.8; exp: 1.1±1.1 Hz) and movement (original model: 4.0±5.7 Hz; modified model for MTh inactivation: 3.0 ± 5.0; exp: 2.2 ± 4.0 Hz) states, supporting our hypothesis of the circuitry involved in the MTh inactivation condition.

### M1 dynamics during noradrenergic (NA) receptor blockade

We then explored the role of NA neuromodulation in the model, motivated by our in vivo study where blocking NA inputs through local infusion of NA-R antagonists resulted in reduced motor coordination (***Schiemann et al., 2015***). Other studies have also shown that NA alters M1 signaling during movement and motor behavior (***Dacre et al., 2021; Guo et al., 2021; Sheets et al., 2011**).* The model reproduced key aspects of the experimental M1 L5B responses under this noradrenergic receptor blockade (NA-R block) condition. NA-R block was initially simulated by resetting *I*_h_ from the in vivo to the baseline in vitro condition (100% *I*_h_ conductance in PT cells), reflecting no NA input from LC. Long-range inputs from seven cortical and thalamic regions were unchanged from the control condition. Under NA-R block condition, the change from quiet to movement states only involved increasing the firing rate of MTh inputs. NA-R block resulted in decreased L5B activation during movement compared to control condition (Fig. 5C,D) (control: 6.9 ± 9.7 Hz, NA-R block: 5.6 ± 6.2 Hz), particularly in the PT5B population (control: 11.9 ± 11.3 Hz, NA-R block: 5.1 ± 6.3 Hz). In vivo experiments showed a more pronounced decrease in L5B movement rates (control: 8.4 ± 7.5 Hz, NA-R block: 1.3 ± 2.2 Hz). A similar decrease during NA-R block was observed in the quiet rates of L5B and L5B IT, whereas these model populations remained at a similar rate than in the control condition.

These discrepancies between experiment and model suggested that the model was not fully capturing some effects of noradrenergic LC inputs. As in the MTh inactivation condition, this provided an opportunity to evaluate hypotheses that could improve future versions of the model. We therefore tested one possible hypothesis by modifying the model to incorporate an additional known effect of NA, namely, the modulation of potassium (*K*^+^) conductance ***(Wang and McCormick, 1993; Favero et al., 2012; Schiemann et al., 2015***). Increased NA has been shown to reduce *K^+^* conductance, hence to simulate this effect during the NA-block condition we increased potassium conductance by 50% in all excitatory cell types. The combined effect of increasing *I*_h_ and *K^+^* better captured the experimental responses during the NA-block condition (see Fig. 5D purple lines). More specifically, L5B, L5 IT and L5B_suppressed_ mean firing rates were lower for both the quiet (L5B IT: orig model: 7.0 ± 6.1; modified model for NA-R block: 1.7 ± 2.6; exp: 2.0 ± 0.7 Hz) and movement (L5B IT: orig model: 6.2 ± 6.0; modified model for NA-R block: 1.1 ± 1.9; exp: 2.4 ± 3.0 Hz) responses, more closely matching those recorded in vivo. This supports the hypothesis that changes in *K^+^* conductance are an important component of LC-mediated NA modulation.

### Motor thalamic and noradrenergic inputs affect L5B dynamics in a cell type and sublayer-specific manner

Our model reproduced the pattern of M1 L5B in vivo responses observed experimentally for different levels of MTh and NA inputs, and provided insights and predictions of how the different L5B subpopulations respond and interact (Fig. 6). The experimental and modeling results reported so farsuggestthat M1 L5B response depends strongly on MTh and NA inputs. Fig. 6A shows the experiment (top) and model (bottom) L5B mean firing rates as a function of these two inputs, illustrating that MTh and NA inputs moderately increased the L5B response, but both are simultaneously re-quired to trigger high L5B activity. Both experiment and model exhibit a similar response pattern, progressively increasing with MTh and NA, and a similar range of L5B firing rates. We note that these experimental results combine and extrapolate data from the control, MTh inactivation and NA-R block conditions. The model results corresponds to the original version (without the modifications proposed in the previous sections) but we included additional simulations covering the full parameter space explored, i.e. all combinations of MTh input and NA modulation (PT *I*_h_) values (see Methods for details). To provide a better intuition of the full circuit model dynamics, we also included the spiking raster plots for the 4 conditions with minimum and maximum MTh/NA values (see arrows from the 4 corners of the model heatmap in Fig. 6A).

**Figure 6.**
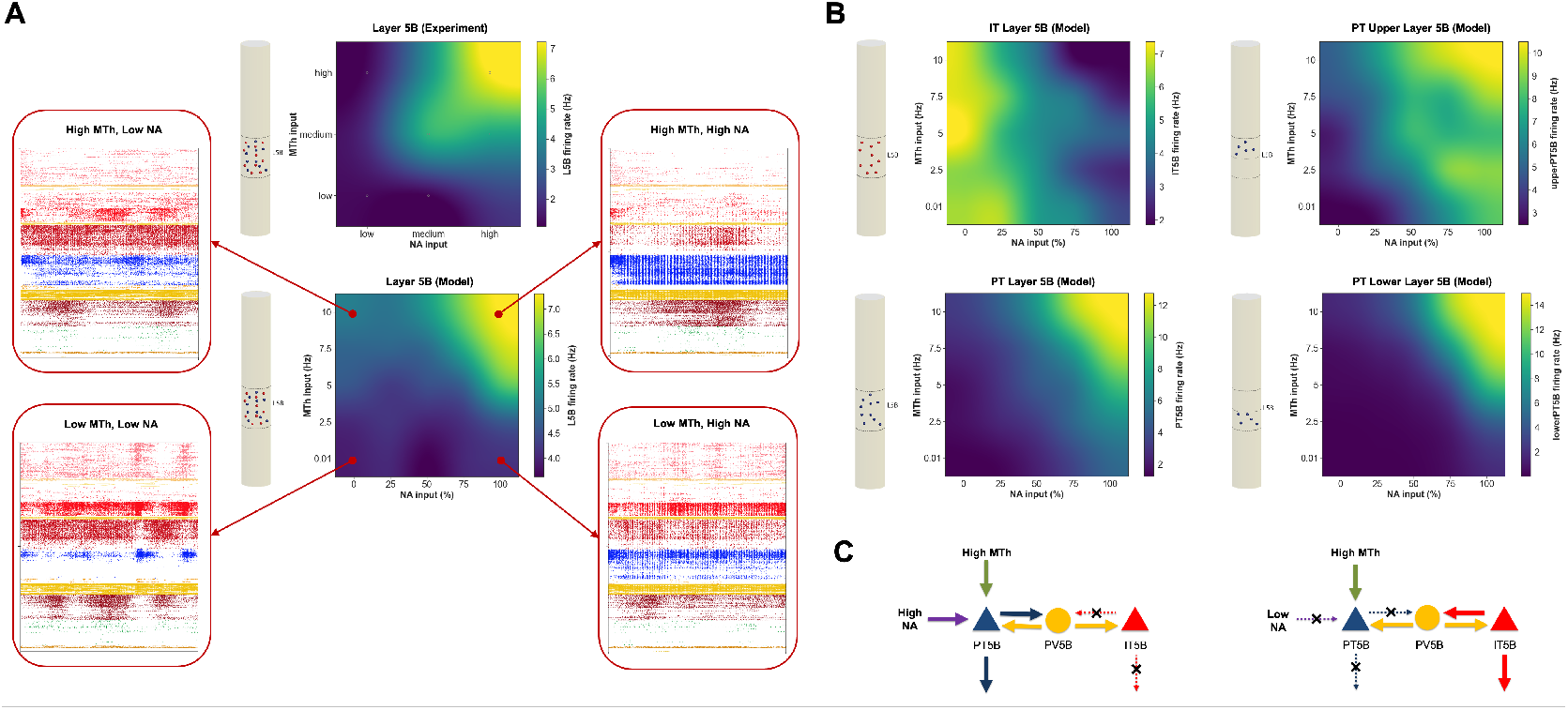
Cell type and sublayer-specific effects of MTh and NA input levels on L5B dynamics. **A.** Mean L5B firing rate response of experiment (top) and model (bottom) to different levels of MTh and NA inputs. Firing raster plot of full circuit model (4 secs) shown inset for each of the four extreme conditions. Schematic cylinders illustrate the cell type (IT=red;PT=blue) and layer analyzed. Experimental values derived from the control, MTh inactivation and NA-R block conditions indicated with small gray circle (remaining values were extrapolated) Model results include additional simulations covering the full parameter space explored. **B.** Same as in A but for different L5B cell types and subpopulations (IT, PT, PT5B_upper_ and PT5B_lower_) each of which showed highly specific response patterns to MTh and NA. **C.** Schematic of hypothesized NA inputs and mutual disynaptic inhibitory pathway mediating the switching between IT- and PT-predominant output modes.

The model revealed highly specific and distinct activity patterns for the different L5B cell types and sublayers (Fig. 6B). Somewhat surprisingly, L5B IT cells exhibited an inverse response pattern to NA compared to L5B PT and to the overall L5B response (Fig. 6B), showing decreased firing with increases of NA inputs;and a largely constant response to MTh inputs. The NA response is consistent with the low levels of *I*_h_ expression in L5B IT cells (***Sheets et al., 2011***). We hypothesize the inverse response to NA between L5B IT and PT cells could be caused by mutual inhibition mediated via L5 interneurons (see schematic in Fig. 6C). L5B PT cells showed higher peak firing rates than IT (12.8 Hz vs 7.4 Hz) thus dictating the overall L5B response pattern and overshadowing L5 IT inverse pattern. Supragranular IT2/3 and IT5A populations exhibited generally low activity (see Fig. 6A raster plots) when PT5B fired strongly (high MTh and NA), consistent with the predominant involvement of PT cells in motor execution (***Li et al., 2015b**).* The model also exposed sublaminar differences in L5B PT response, with PT5B_lower_ exhibiting more extreme minimum and maximum rates than PT5B_upper_ (0 – 15 Hz vs 3 – 10 Hz). The PT5B_lower_ activation threshold was also higher than for PT5B_upper_, i.e. PT5B_upper_ required higher MTh and NA inputs to start responding strongly. This suggests PT5B_upper_ would activate first followed by a delayed response from PT5B_lower_, as inputs associated with motor execution accumulate and reach a threshold. These results in line with the suggested role of PT5B_upper_ in movement preparation and PT5B_lower_ cells in movement initiation (***Economo et al., 2018***).

## Discussion

In this work, we have developed a computational model of the mouse M1 microcircuit and validated it against in vivo data. Despite inherent limitations due to gaps in the data (see details in the section below), we believe that this constitutes the most biophysically detailed model of mouse M1 currently available comprising the molecular, cellular and circuit scales. The model integrates quantitative experimental data on neuronal physiology, morphology, laminar density, cell type distribution, dendritic distribution of synapses, and local and long-range synaptic connectivity, obtained from 31 studies, with 12 of these coming from our experimental laboratory. Model development also benefited greatly from extended discussions between the computational and experimental authors. Integrating data across scales and managing such a complex model motivated the development of a novel software tool, NetPyNE, that provides a high-level interface to NEURON and facilitates multiscale brain circuit modeling (***Dura-Bernal et al., 2019***).

To validate the model we focused on reproducing mouse M1 in vivo experimental results across different behavioral states and experimental conditions from a single study ***(Schiemann et al., 2015)**.* Simulation results were consistent across multiple random wiring seeds and background input seeds, demonstrating the robustness of the model. The model cell type-specific spontaneous firing rates, associated with the quiet behavior, were consistent with experimental data from several in vivo studies (***Schiemann et al., 2015; Zagha et al., 2015; Li et al., 2016; Estebanez et al., 2018; Economo et al., 2018**)* (Fig. 2). We then simulated activity corresponding to mouse self-paced, voluntary locomotion by increasing motor thalamus (MTh) and noradrenaline (NA) inputs. Movement-related changes in L2/3 and L5B population firing rates were consistent with those reported in vivo, including bidirectional (enhanced vs suppressed) firing rate changes in distinct L5B pyramidal neuron populations (Fig. 3). Local field potentials (LFP) oscillations emerged spontaneously (no oscillatory inputs) at physiological frequencies, and included characteristic delta, beta and gamma oscillatory patterns. LFP power in L5B shifted from lower (delta) to higher (gamma) frequency bands during movement, consistent with in vivo LFP data (Fig. 4).

We also simulated two experimental manipulations – inactivation of MTh inputs and blocking of NA receptors – which resulted in cell type-specific activity changes in L5B which matched those measured experimentally (Fig. 5). For each condition, we evaluated two hypotheses of the cellular and circuit mechanisms involved, which suggested MTh inactivation may affect other long-range inputs, and NA modulation affects not only *I*_h_ but also K+ conductances. We used the model to systematically explore the interaction between MTh and NA inputs and predict M1 output at the level of individual cell types at sublaminar resolution. Results captured the overall pattern and response amplitudes measured in vivo, supporting the hypotheses both high MTh and NA inputs are required for self-paced voluntary movement-related L5B activity (Fig. 6). The model predicted a predominant role of PT cells in dictating L5B responses during movement, with PT5B_lower_ providing the strongest response but only when both MTh and NA inputs were high enough, i.e. PT5B_lower_ exhibited the highest response threshold. L5B IT cells exhibited an opposite but lower-amplitude pattern, potentially due to PT-mediated disynaptic inhibition, and infragranular IT were less engaged during the movement state. These predictions are consistent with findings associating IT and PT5B_upper_ with motor planning and PT5B_lower_ with motor execution (***Li et al., 2015b; Economo et al., 2018)**.*

This is, to the best of our knowledge, the first model of the mouse M1 microcircuit where firing rates and LFPs have been directly compared to cell type and layer-specific mouse M1 in vivo data associated with different behaviors and experimental manipulations. The model provides a quantitative theoretical framework to integrate and interpret M1 experimental data across scales, evaluate hypotheses and generate experimentally testable predictions.

### Challenges and limitations

Our ambition was to develop a detailed multiscale computational model of the mouse M1 microcircuit. We necessarily fell short due to lack of data of some molecular, cellular, network and long-range connectivity details. This model was constructed and evaluated over a period of six years. During this period we updated the model multiple times to incorporate new data, but of course any neurobiological model is always in need of additional updating and improvement as new measurements become available.

Of some concern is the relative lack of data on dendritic ion channel density, which will affect the influence of distal synaptic inputs on L5 neurons (***Labarrera et al., 2018**).* Cell models are precisely tuned to reproduce experimental somatic responses, but limited data is available to characterize dendritic physiology. Although we adapted the morphology and physiology of IT cells based on their layer, we omitted cellular diversity within each model population – all the model neurons of the same cell type and layer have identical morphologies and identical channel parameters. This contrasts with other models which vary both channel conductances and morphologies, the latter by slightly jittering angles and lengths (***Markram et al., 2015a**).*

Due to the nature of our circuit mapping methods (***Anderson et al., 2010; Hooks et al., 2013; Suter and Shepherd, 2015***), our model used local excitatory connectivity primarily based on post-synaptic cell type and presynaptic locations. Our model’s normalized cortical-depth-dependent connectivity provided greater resolution than traditional layer-based wiring, but still contained boundaries where connection density changed and did not provide cell level point-to-point resolution. This could be further improved by fitting discretely binned experimental data to functions of cortical depth, resulting in smoother connectivity profiles. Other recent models have used a sophisticated version of Peters’ principle (identifying overlap between axonal and dendritic trees) to provide cell-to-cell resolution for selected cells, which must then still be replicated and generalized across multiple instances to build a large network (***Rees et al., 2017; Markram et al., 2015a***). Inclusion of synpatic plasticity mechanisms could be used to study the role of different cell types in motor learning, for example, L5A neurons which evidence suggests participate in the evolving network representation of learned movements (***Masamizu et al., 2014***).

We are limited not only by lack of precise data for parameter determination, but also by computational constraints. Often, network simulations use point neurons in order to avoid the computational load of multicompartment neurons, but at the expense of accuracy ***(Potjans and Diesmann, 2014; Izhikevich and Edelman, 2008; Schmidt et al., 2018***). Here, we compromised by using relatively small multicompartment models for most populations, with the exception of the neurons of L5. In terms of noradrenaline influence, we focused here on one effect on the PT cell type, neglecting the wide-ranging effects of this and other neuromodulators (such as dopamine, acetylcholine) (***O’Donnell et al., 2012; McCormick, 1992; Graybiel, 1990**)* and their the influence of second messenger cascades (***Neymotin et al., 2016a***). Implementing this functionality is now available via NEURON’s *rxd* module (***McDougal et al., 2013; Newton et al., 2018***). Even with these compromises, optimizing and exploring our large network model required millions of HPC core-hours.

In summary, model firing rate distributions were generally consistent with experimental data across populations and behavioral states. We note that the experimental dataset represents a small sparse sample of neurons in the modeled cortical volume, resulting in a model data sample size approximately 3 orders of magnitude larger than that of experiment (e.g. for L5B *N_model_* = 35182 vs *N_experiment_* = 47). Therefore, validation of our model results can be understood as showing that the small dataset of experiment cell rates could have been subsampled from the larger dataset of model rates. Novel methods that record from an increasingly larger number of simultaneous neurons (***Hong and Lieber, 2019**)* will enable further validation of the model results.

### M1 cellular and circuit mechanisms associated with quiet and movement behaviors

A key question in motor system research is how motor cortex activity gets dissociated from muscle movement during motor planning or mental imagery, and is then shifted to produce commands for action (***Ebbesen and Brecht, 2017; Schieber, 2011; Shenoy et al., 2013***). One hypothesis has been that this planning-to-execution switch might be triggered by NA neuromodulation (***Sheets et al., 2011***). Downregulation of *I*_h_, effected via NA and other neuromodulatory factors, has been shown to increase PT activity as a consequence of enhanced temporal and spatial synaptic integration of EPSPs (***Sheets et al., 2011; Labarrera et al., 2018**).* This effect is primarily observed in PT cells, since the concentration of HCN channels in these cells has been shown to be significantly higher than in IT cells (***Sheets et al., 2011; BICCN, 2021***). In the model, we used a baseline *I*_h_ consistent with a cell tuned to reproduce in vitro data with no NA modulation. For the in vivo quiet condition (low NA modulation), we used 75% of that baseline level, and for movement (high NA) we used 25%, consistent with values reported experimentally (***Labarrera et al., 2018***). Paradoxically, *I*_h_ downregulation has also been reported to *reduce* pyramidal cell activity in some settings (***George et al., 2009; Migliore and Migliore, 2012***). Here we improved our previous PT cell model (***Neymotin et al., 2017**)* to include an *I*_h_ model (***Migliore and Migliore, 2012**)* that was able to reconcile these observations: *I*_h_ downregulation reduced PT response to weak inputs, while increasing the cell response to strong inputs (***Migliore and Migliore, 2012; George et al., 2009; Sheets et al., 2011; Labarrera et al., 2018**).*

An additional hypothesis to explain differential planning and movement outputs, posits that the shift results from activation of different cell populations in L5, mediated by distinct local and long-range inputs. Accumulated evidence suggests that inputs arising from MTh carrying cerebellar signals differentially target M1 populations (***Hooks et al., 2013***) and are involved in triggering movement (***Dacre et al., 2021**)* and in dexterous tasks (***Guo et al., 2021***). Using in vivo electrophysiology and optogenetic perturbations in mouse anterolateral motor cortex, ***Li et al. (2015b)*** found evidence suggesting that preparatory activity in IT neurons is converted into a movement command in PT neurons. Further support for this hypothesis comes from a study that showed that transcriptomically-identified different PT subtypes in upper vs lower L5B (***Economo et al., 2018**),* and showed that PT5B_upper_ projected to thalamus and generated early preparatory activity, while PT5B_lower_ projected to medulla and generated motor commands.

These two hypotheses are not incompatible, and indeed our simulations suggest that both of these mechanisms may coexist and be required for movement-related activity (Fig. 6). NA modulation and MTh input by themselves produced an increase in PT5B overall activity, but primarily in the preparatory activity-related PT5B_upper_ population; both mechanisms were required to activate the PT5B_lower_ population associated with motor commands (***Economo et al., 2018**).* The model therefore predicts that the transition to motor execution (self-paced, voluntary movement) might require both the neuromodulatory prepared state and circuit-level routing of inputs. Different types of behaviors and contexts (e.g. goal-directed behaviors with sensory feedback) may involve driving inputs from other populations or regions, such as supragranular layers or somatosensory cortex (***Hooks et al., 2013; Dacre et al., 2021; Zareian et al., 2021; Yamawaki et al., 2021**).* We note that in our model and in vivo experiments (***Schiemann et al., 2015**)* the quiet state does not correspond to a preparatory state, as it lacks short-term memory, delays and other preparatory components. Therefore, whether previous task-related findings (***Li et al., 2015b; Economo et al., 2018)*** on the role of PT5B_lower_ and PT5B_upper_ generalize to our self-paced voluntary movement results remains an open question.

### Simulating experimental manipulations: motor thalamus inactivation and noradrenergic receptor blocking

Attempting to reproduce the extreme conditions posed by experimental manipulations provided further insights into the circuitry and mechanisms governing M1 dynamics. During MTh inactivation, our baseline model exhibited higher firing rates than in vivo, particularly for the quiet state. We hypothesized this may be due to inactivation of MTh also affecting other afferent regions of M1, such as contralateral M1 and S2; either directly (e.g. VL→S2) and/or indirectly via recurrent interareal projections (e.g. M1 →S2→M1). We evaluated this by reducing activity in these model regions, which indeed resulted in a closer match to in vivo rates (Fig. 5). Several other hypotheses may also explain the observed discrepancies, for example, that movement-related activity 1) depends on changes in spiking patterns and not just amplitude (e.g. bursts or oscillatory activity); or 2) that it is driven not only by VL but by other long-range inputs (consistent with recent findings ***(Dacre et al., 2021)**),* and/or by local lateral inputs from non-modeled regions of M1. The inclusion of detailed interactions among afferent cortical and thalamic regions is out of the scope of this paper. However, our results already suggested possible improvements to the model and circuit pathways to explore experimentally, demonstrating that the model can be used to evaluate different candidate circuitries and activity patterns.

Similarly, for the NA receptor block condition, we modified the model to evaluate the hypothesis that it not only increases PT *I*_h_ but also K+ conductance in all pyramidal neurons, as suggested by multiple studies (***Wang and McCormick, 1993; Favero et al., 2012**).* This resulted in a closer match between model and experiment. Alternative hypotheses that may also account for the initial differences observed include NA selective modulation of inhibitory synapses, and interactions with other neuromodulators such as acetylcholine (***Conner et al., 2010**).* These molecular and cellular level mechanisms can be explored in our model to gain insights into their circuit-level effects.

### IT and PT disynaptic inhibition via shared L5 interneuron pools

L5B IT and PT neurons exhibited an inverse response to increased NA inputs: IT rates decreased while PT rates increased (Fig. 6B). We hypothesize this effect may result from mutual inhibition between IT and PT mediated via a shared pool of L5 interneurons, as illustrated in the schematic in (Fig. 6C). This is in line with the finding of shared interneuron pools in L5 IT and PT neurons mediating disynaptic inhibition (***Apicella et al., 2012***), which contrast with the private (non-shared) interneuron pools identified for PT and CT neurons (***Yamawaki and Shepherd, 2015**).* Additional support comes from rat in vivo results showing PV neurons were recruited predominantly during motor execution and may shape motor commands through balanced or recurrent inhibition of output-related pyramidal neurons (PT), while supressing pyramidal neurons (IT) associated with other functions such as hold-related activity (***Isomura et al., 2009**).* By modeling the M1 circuit connectivity and simulating its dynamics we have predicted the computation performed by this particular subcircuit, namely, a switching mechanism between IT- and PT-predominant output modes (mutual inhibition ensures only one of them responds strongly at a time). This is consistent with their suggested complementary roles in motor preparation vs execution (***Li et al., 2015b**).* This circuit-level prediction can be tested experimentally in future studies.

### Emergence of behavior-dependent physiological oscillations

Our model of M1 neocortex exhibits spontaneous physiological oscillations without rhythmogenic synaptic input. Strong oscillations were observed in the delta and beta/gamma ranges with specific frequency-dependence on cell class, cortical depth, and behavioral state. The simulated reproduced the decrease in delta and increase in gamma power of M1 L5 LFP during movement observed in the in vivo dataset (***Schiemann et al., 2015***), and previously reported in mouse vibrissal M1 during whisking (***Zagha et al., 2013***). The model can be used to provide cell type-specific predictions as to the origins of behavior-related changes in LFP. For example, given the increase of PT firing rates and oscillatory activity observed during movement (Fig. 3), we hypothesized that the movement-related increase in L5 LFP gamma oscillations is largely mediated by PT neurons. Strong LFP beta and gamma oscillations are characteristic of motor cortex activity in both rodents (***Castro-Alamancos, 2013; Tsubo et al., 2013**)* and primates (***Rubino et al., 2006; Nishimura et al., 2013***), and have been found to enhance signal transmission in mouse neocortex (***Sohal et al., 2009**).* Both beta and gamma oscillations may play a role in information coding during preparation and execution of movements (***Ainsworth et al., 2012; Tsubo et al., 2013***). More generally, these physiological oscillations are considered to be fundamental to the relation of brain structure and function (***Buzsáki and Mizuseki, 2014***). As the primary output, PT cells receive and integrate many local and long-range inputs. Their only local connections to other L5 excitatory neurons are to other PT cells (***Kiritani et al., 2012***). However, as described in the previous section, by targeting inhibitory cells in L5 they are able to reach across layers to influence other excitatory populations, either reducing activity or entraining activity (***Naka and Adesnik, 2016***). These disynaptic E→I→E pathways likely play a role in coupling oscillations within and across layers, and in setting frequency bands.

### Implications for experimental research and therapeutics

Our model integrates previously isolated experimental data at multiple scales into a unified simulation that can be progressively extended as new data become available. This provides a useful tool for researchers in the field, who can use this quantitative theoretical framework to evaluate hypotheses, make predictions and guide the design of new experiments using our freely-available model (see Methods). This in silico testbed can be systematically probed to study microcircuit dynamics and biophysical mechanisms with a level of resolution and precision not available experimentally. Unraveling the non-intuitive multiscale interactions occurring in M1 circuits can help us understand disease mechanisms and develop new pharmacological and neurostimulation treatments for brain disorders (***Neymotin et al., 2016c,b; Dura-Bernal et al., 2016; Arle and Shils, 2008; Wang et al., 2015; Bensmaia and Miller, 2014; Sanchez et al., 2012***), and improve decoding methods for brain-machine interfaces (***Carmena, 2013; Shenoy and Carmena, 2014; Dura-Bernal et al., 2017; Kocaturk et al., 2015***).

## Methods

The methods below describe model development with data provenance, and major aspects of the final model. The full documentation of the final model is the source code itself, available for download at http://modeldb.yale.edu/260015.

### Morphology and physiology of neuron classes

Seven excitatory pyramidal cell and two interneuron cell models were employed in the network. Their morphology and physiological responses are summarized in Figs. 1*A,B,C* and 7. In previous work we developed layer 5B PT corticospinal cell and L5 IT corticostriatal cell models that reproduced in vitro electrophysiological responses to somatic current injections, including sub- and super-threshold voltage trajectories and f-I curves (***Neymotin et al., 2017; Suter et al., 2013**).* To achieve this, we optimized the parameters of the Hodgkin-Huxley neuron model ionic channels – Na, Kdr, Ka, Kd, HCN, CaL, CaN, KCa – within a range of values constrained by the literature. The corticospinal and corticostriatal cell model morphologies had 706 and 325 compartments, respectively, digitally reconstructed from 3D microscopy images. Morphologies are available via Neuro-Morpho.org (***Ascoli et al., 2007**)* (archive name “Suter_Shepherd”). For the current simulations, we further improved the PT model by 1) increasing the concentration of Ca2+ channels (“hot zones”) between the nexus and apical tuft, following parameters published in (***Hay et al., 2011***); 2) lowering dendritic Na+ channel density in order to increase the threshold required to elicit dendritic spikes, which then required adapting the axon sodium conductance and axial resistance to maintain a similar f-I curve; 3) replacing the HCN channel model and distribution with a more recent implementation (***Migliore and Migliore, 2012***). The new HCN channel reproduced a wider range of experimental observations than our previous implementation (***Kole et al., 2006**),* including the change from excitatory to inhibitory effect in response to synaptic inputs of increasing strength (***George et al., 2009**).* This was achieved by including a shunting current proportional to *I*_h_. We tuned the HCN parameters (*Ik* and *v_re_l_k_*) and passive parameters to reproduce the findings noted above, while keeping a consistent f-I curve consistent (***Suter et al., 2013**).*

The network model includes five other excitatory cell classes: layer 2/3, layer 4, layer 5B and layer 6 IT neurons and layer 6 CT neurons. Since our focus was on the role of L5 neurons, other cell classes were implemented using simpler models as a trade-off to enable running a larger number of exploratory network simulations. Previously we had optimized 6-compartment neuron models to reproduce somatic current clamp recordings from two IT cells in layers 5A and 5B. The layer 5A cell had a lower f-I slope (77 Hz/nA) and higher rheobase (250 nA) than that in layer 5B (98 Hz/nA and 100 nA). Based on our own and published data, we found two broad IT categories based on projection and intrinsic properties: corticocortical IT cells found in upper layers 2/3 and 4 which exhibited a lower f-I slope (~72 Hz/nA) and higher rheobase (~281 pA) than IT corticostriatal cells in deeper layers 5A, 5B and 6 (~96 Hz/nA and ~106 pA) (***Yamawaki et al., 2015; Suter et al., 2013; Oswald et al., 2013***). CT neurons’ f-I rheobase and slope (69 Hz/nA and 298 pA) was closer to that of corticocortical neurons (***Oswald et al., 2013**).* We therefore employed the layer 5A IT model for layers 2/3 and 4 IT neurons and layer 6 CT neurons, and the layer 5B IT model for layers 5A, 5B and 6 IT neurons. We further adapted cell models by modifying their apical dendrite length to match the average cortical depth of the layer, thus introducing small variations in the firing responses of neurons across layers.

We implemented models for two major classes of GABAergic interneurons (***Huang, 2014; BICCN, 2021; Rudy et al., 2011**):* parvalbumin-expressing fast-spiking (PV) and somatostatin-expressing low-threshold spiking neurons (SOM). We employed existing simplified 3-compartment (soma, axon, dendrite) models (***Konstantoudaki et al., 2014**)* and increased their dendritic length to better match the average f-I slope and rheobase experimental values of cortical basket (PV) and Martinotti (SOM) cells (Neuroelectro online database (***Tripathy et al., 2015**)).*

### Microcircuit composition: neuron locations, densities and ratios

We modeled a cylindric volume of the mouse M1 cortical microcircuit with a 300 *μm* diameter and 1350 *μm* height (cortical depth) at full neuronal density for a total of 10,073 neurons (Fig. 1). Cylinder diameter was chosen to approximately match the horizontal dendritic span of a corticospinal neuron located at the center, consistent with the approach used in the Human Brain Project model of the rat S1 microcircuit (***Markram et al., 2015b***). Mouse cortical depth and boundaries for layers 2/3, 4, 5A, 5B and 6 were based on our published experimental data (***Weiler et al., 2008; Anderson et al., 2010; Yamawaki et al., 2015***). Although traditionally M1 has been considered an agranular area lacking layer 4, we recently identified M1 pyramidal neurons with the expected prototypical physiological, morphological and wiring properties of layer 4 neurons (***Yamawaki et al., 2015***) (see also (***Bopp et al., 2017; Barbas and García-Cabezas, 2015; BICCN, 2021***)), and therefore incorporated this layer in the model.

Cell classes present in each layer were determined based on mouse M1 studies ***(Suter et al., 2013; Anderson et al., 2010; Yamawaki et al., 2015; Oswald et al., 2013; Naka and Adesnik, 2016***). IT cell populations were present in all layers, whereas the PT cell population was confined to layer 5B, and the CT cell population only occupied layer 6. SOM and PV interneuron populations were distributed in each layer. Neuronal densities (neurons per *mm*^3^) for each layer (Fig. 1*C*) were taken from a histological and imaging study of mouse agranular cortex (***Tsai et al., 2009***). The proportion of excitatory to inhibitory neurons per layer was obtained from mouse S1 data (***Lefort et al., 2009***). The proportion of IT to PT and IT to CT cells in layers 5B and 6, respectively, were both estimated as 1:1 (***Suter et al., 2013; Yamawaki and Shepherd, 2015***). The ratio of PV to SOM neurons per layer was estimated as 2:1 based on mouse M1 and S1 studies (***Katzel et al., 2011; Wall et al., 2016***) (Fig. 7*B*). Since data for M1 layer 4 was not available, interneuron populations labeled PV5A and SOM5A occupy both layers 4 and 5A. The number of cells for each population was calculated based on the modeled cylinder dimensions, layer boundaries and neuronal proportions and densities per layer.

**Figure 7.**
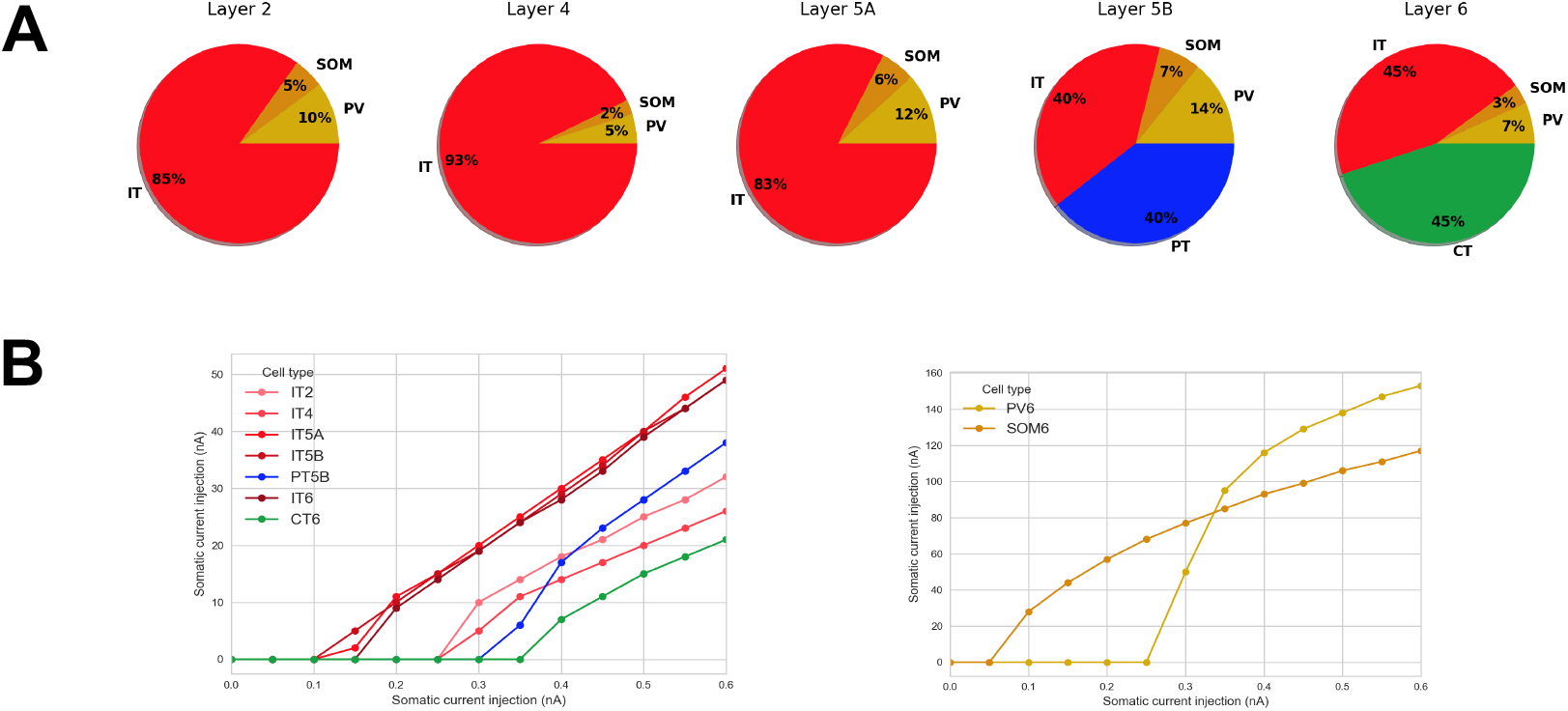
Microcircuit layer composition and cell type f-I response. **A.** Proportion of cell classes per layer; **B.** f-I curve for each excitatory and inhibitory cell types. All properties were derived from published experimental data. Populations labels include the cell class and layer, e.g. ‘IT2’ represents the IT class neurons in layer 2/3.

### Local connectivity

We calculated local connectivity between M1 neurons (Figures *1C* and 8*A*) by combining data from multiple studies. Data on excitatory inputs to excitatory neurons (IT, PT and CT) was primarily derived from mapping studies using whole-cell recording, glutamate uncaging-based laserscanning photostimulation (LSPS) and subcellular channelrhodopsin-2-assisted circuit mapping (sCRACM) analysis (***Weiler et al., 2008; Anderson et al., 2010; Yamawaki et al., 2015; Yamawaki and Shepherd, 2015)**.* Connectivity data was postsynaptic cell class-specific and employed normalized cortical depth (NCD) instead of layers as the primary reference system. Unlike layer definitions which can be interpreted differently between studies, NCD provides a well-defined, consistent and continuous reference system, depending only on two readily-identifiable landmarks: pia (NCD=0) and white matter (NCD=1). Incorporating NCD-based connectivity into our model allowed us to capture wiring patterns down to a 100 *μm* spatial resolution, well beyond traditional layer-based cortical models. M1 connectivity varied systematically within layers. For example, the strength of inputs from layer 2/3 to L5B corticospinal cells depends significantly on cell soma depth, with upper neurons receiving much stronger input (***Anderson et al., 2010**).*

**Figure 8.**
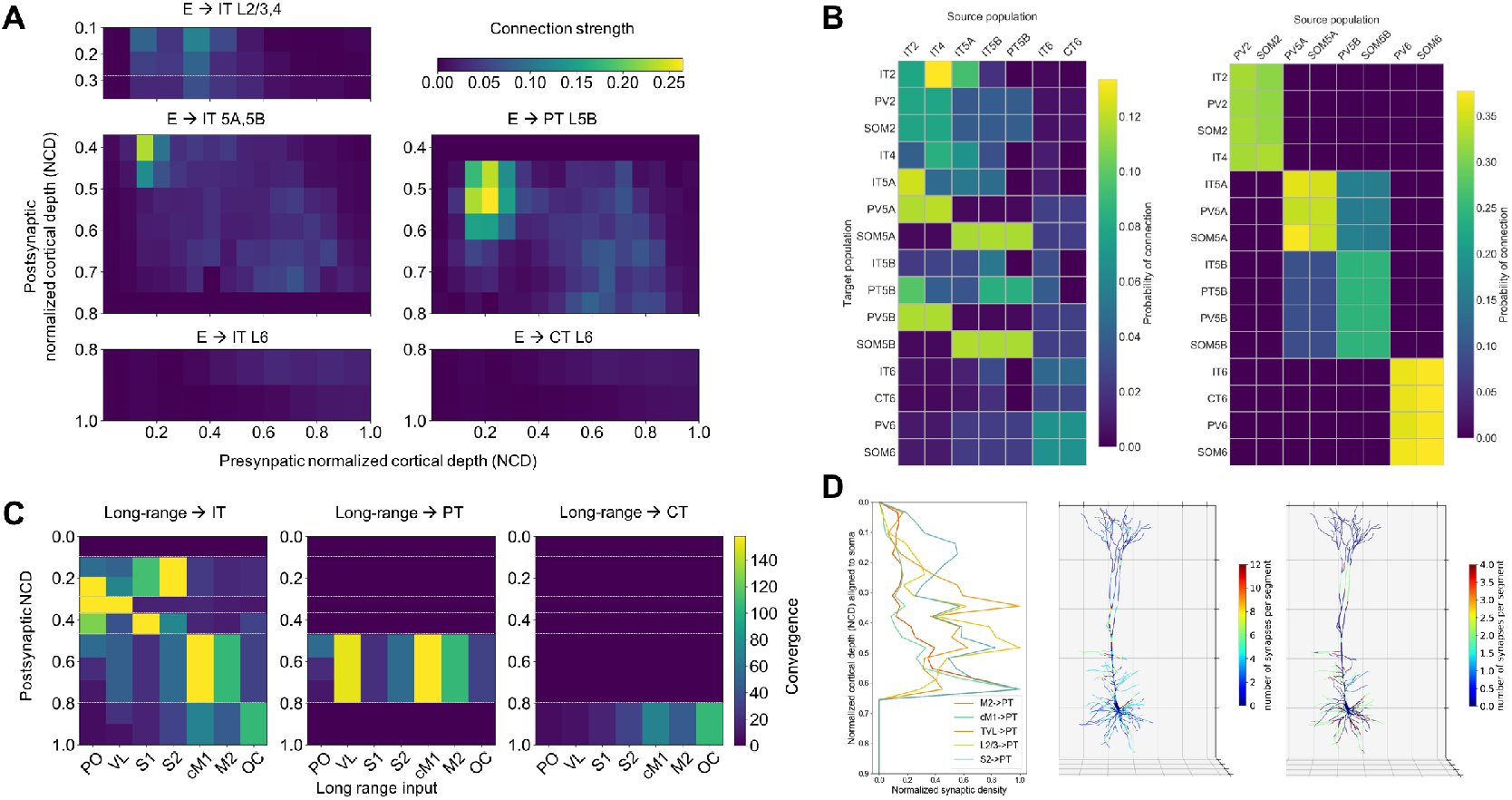
M1 excitatory connectivity: local microcircuitry and and long-range inputs. **A.** Strength of local excitatory connections as a function of pre- and post-synaptic normalized cortical depth (NCD) and post-synaptic cell class;values used to construct the network. **B.** Convergence of long-range excitatory inputs from seven thalamic and cortical regions as a function post-synaptic NCD and cell class;values used to construct the network. **C.** Probability of connection matrix for excitatory (left) and inhibitory (right) populations calculated from an instantiation of the base model network. **D.** Left. Synaptic density profile (1D) along the dendritic arbor for inputs from layer 2/3 IT, VL, S1, S2, cM1 and M2 to PT neurons. Calculated by normalizing sCRACM maps (*(**Suter and Shepherd, 2015**)* Figs. 5 and 6) by dendritic length at each grid location and averaging across rows. Middle and Right. Synaptic density per neuron segment automatically calculated for each neuron based on its morphology and the pre- and postsynaptic cell type-specific radial synaptic density function. Here, VL →PT and S2 →PT are compared and exhibit partially complementary distributions.

Connection strength thus depended on presynaptic NCD and postsynaptic NCD and cell class. For postsynaptic IT neurons with NCD ranging from 0.1 to 0.37 (layers 2/3 and 4) and 0.8 to 1.0 (layer 6) we determined connection strengths based on data from (***Weiler et al., 2008***) with cortical depth resolution of 140 *μm*-resolution. For postsynaptic IT and PT neurons with NCD between 0.37 and 0.8 (layers 5Aand 5B) we employed connectivity strength data from (***Anderson et al., 2010***) with cortical depth resolution of 100 *μm.* For postsynaptic CT neurons in layer 6 we used the same connection strengths as for layer 6 IT cells (***Weiler et al., 2008**),* but reduced to 62% of original values, following published data on the circuitry of M1 CT neurons (***Yamawaki and Shepherd, 2015; Kuramoto et al., 2022***). Our data (***Yamawaki and Shepherd, 2015***) also suggested that connection strength from layer 4 to layer 2/3 IT cells was similar to that measured in S1, so for these projections we employed values from Lefort’s S1 connectivity strength matrix (***Lefort et al., 2009***). Experimentally, these connections were found to be four times stronger than in the opposite direction – from layer 2/3 to layer 4 – so we decreased the latter in the model to match this ratio.

Following previous publications (***Kiritani et al., 2012; Lefort et al., 2009***), we defined connection strength (*s_cmt’_*, in mV) between two populations, as the product of their probability of connection (*p_cm_*) and the unitary connection somatic EPSP amplitude in mV (*v_con_*), i.e. *s_con_* = *p_con_* × *v_con_*. We employed this equivalence to disentangle the connection *s_con_* values provided by the above LSPS studies into *p_con_* and *v_con_* values that we could use to implement the model. First, we rescaled the LSPS raw current values in pA (***Anderson et al., 2010; Weiler et al., 2008; Yamawaki et al., 2015; Yamawaki and Shepherd, 2015***) to match *s_con_* data from a paired recording study of mouse M1 L5 excitatory circuits (***Kiritani et al., 2012**).* Next, we calculated the M1 NCD-based *v_con_* matrix by interpolating a layerwise unitary connection EPSP amplitude matrix of mouse S1 ***(Lefort et al., 2009***), and thresholding values between 0.3 and 1.0 mV. Finally, we calculated the probability of connection matrix as *P_con_* = *s_con_/V_cmt_*.

To implement *v_con_* values in the model we calculated the required NEURON connection weight of an excitatory synaptic input to generate a somatic EPSP of 0.5 mV at each neuron segment. This allowed us to calculate a scaling factor for each segment that converted *v_com_* values into NEURON weights, such that the somatic EPSP response to a unitary connection input was independent of synaptic location – also known as synaptic democracy (***Rumsey and Abbott, 2006; Poirazi and Papoutsi, 2020***). This is consistent with experimental evidence showing synaptic conductances increased with distance from soma, to normalize somatic EPSP amplitude of inputs within 300 *μm* of soma (***Magee and Cook, 2000***). Following this study, scaling factor values above 4.0 – such as those calculated for PT cell apical tufts – were thresholded to avoid overexcitability in the network context where each cell receives hundreds of inputs that interact nonlinearly (***Spruston, 2008; Behabadi et al., 2012***). For morphologically detailed cells (layer 5A IT and layer 5B PT), the number of synaptic contacts per unitary connection (or simply, synapses per connection) was set to five, an estimated average consistent with the limited mouse M1 data (***Hu and Agmon, 2016***) and rat S1 studies (***Bruno and Sakmann, 2006; Markram et al., 2015b***). I ndividual synaptic weights were calculated by dividing the unitary connection weight (*v_con_*) by the number of synapses per connection. Although the method does not account for nonlinear summation effects (***Spruston, 2008***), it provides a reasonable approximation and enables employing a more realistic number and spatial distribution of synapses, which maybe key for dendritic computations (***London and Häusser, 2005***). For the remaining cell models, all with six compartments or less, a single synapse per connection was used.

For excitatory inputs to inhibitory cell types (PV and SOM) we started with the same values as for IT cell types but adapted these based on the specific connectivity patterns reported for mouse M1 interneurons (***Apicella et al., 2012; Yamawaki and Shepherd, 2015***) (Fig. 8*A*). Following the layer-based description in these studies, we employed three major subdivisions: layer 2/3 (NCD 0.12 to 0.31), layers 4, 5A and 5B (NCD 0.31 to 0.77) and layer 6 (NCD 0.77 to 1.0). We increased the probability of layer 2/3 excitatory connections to layers 4, 5A and 5B SOM cells by 50% and decreased that to PV cells by 50% (***Apicella et al., 2012***). We implemented the opposite pattern for excitatory connections arising from layer 4,5A,5B IT cells such that PV interneurons received stronger intralaminar inputs than SOM cells (***Apicella et al., 2012***). The model also accounts for layer 6 CT neurons generating relatively more inhibition than IT neurons (***Yamawaki and Shepherd, 2015; Kuramoto et al., 2022***). Inhibitory connections from interneurons (PV and SOM) to other cell types were limited to neurons in the same layer (***Katzel et al., 2011***), with layers 4, 5A and 5B combined into a single layer (***Naka and Adesnik, 2016***). Probability of connection decayed exponentially with the distance between the pre- and post-synaptic cell bodies with length constant of 100 *μm* (***Gal et al., 2017; Fino and Yuste, 2011***). We introduced a correction factor to the distance-dependent connectivity measures to avoid the *border effect,* i.e. cells near the modeled volume edges receiving less or weaker connections than those in the center.

For comparison with other models and experiments, we calculated the probability of connection matrices arranged by population (instead of NCD) for the base model network instantiation used throughout the results. (Fig. 8*B*).

Excitatory synapses consisted of colocalized AMPA(rise, decay *τ*: 0.05, 5.3 ms) and NMDA(rise, decay *τ*: 15,150 ms) receptors, both with reversal potential of 0 mV. The ratio of NMDAto AMPA receptors was 1.0 (***Myme et al., 2003***), meaning their weights were each set to 50% of the connection weight. NMDA conductance was scaled by 1/(1 + 0.28 · *Mg* · exp (−0.062 · *V*)); Mg = 1mM ***(Jahr and Stevens, 1990b***). Inhibitory synapses from SOM to excitatory neurons consisted of a slow *GABA_A_* receptor (rise, decay *τ*: 2,100 ms) and *GABA_B_* receptor, in a 90% to 10% proportion; synapses from SOM to inhibitory neurons only included the slow *GABA_A_* receptor; and synapses from PV to other neurons consisted of a fast *GABA_A_* receptor (rise, decay *τ*: 0.07,18.2). The reversal potential was −80 mV for *GABA_A_* and −95 mV for *GABA_B_*. The *GABA_B_* synapse was modeled using second messenger connectivity to a G protein-coupled inwardly-rectifying potassium channel (GIRK) ***(Destexhe et al., 1996***). The remaining synapses were modeled with a double-exponential mechanism.

Connection delays were estimated as 2 ms plus a variable delay depending on the distance between the pre- and postsynaptic cell bodies assuming a propagation speed of 0.5 m/s.

### Long-range input connectivity

We added long-range input connections from seven regions that are known to project to M1: thalamic posterior nucleus (PO), ventro-lateral thalamus (VL), primary somatosensory cortex (S1), secondary somatosensory cortex (S2), contralateral primary motor cortex (cM1), secondary motor cortex (M2) and orbital cortex (OC). We note that VL constitutes the largest nuclei of the motor thalamus (MTh) so, in the context of the model, these terms are equivalent. Each region consisted of a population of 1000 (***Constantinople and Bruno, 2013; Bruno and Sakmann, 2006***) spike-generators (NEURON VecStims) that generated independent random Poisson spike trains with uniform distributed rates between 0 and 2.5 Hz or 0 and 5 Hz (***Yamashita et al., 2013; Hirata and Castro-Alamancos, 2006***) for spontaneous firing; or 0 and 10 Hz (***Isomura et al., 2009; Jacob et al., 2012***) when simulating increased input from a region. Previous studies provided a measure of normalized input strength from these regions as a function of postsynaptic cell type and layer or NCD. Broadly, PO (***Yamawaki et al., 2015; Yamawaki and Shepherd, 2015; Hooks et al., 2013***), S1 ***(Mao et al., 2011; Yamawaki et al., 2021**)* and S2 (***Suter and Shepherd, 2015***) projected strongly to IT cells in layers 2/3 and 5A (PO also to layer 4); VL projected strongly to PT cells and to layer 4 IT cells (***Yamawaki et al., 2015; Yamawaki and Shepherd, 2015; Hooks et al., 2013***); cM1 and M2 projected strongly to IT and PT cells in layers 5B and 6 (***Hooks et al., 2013**);* and OC projected strongly to layer 6 CT and IT cells (***Hooks et al., 2013**).* We implemented these relations by estimating the maximum number of synaptic inputs from each region and multiplying that value by the normalized input strength for each postsynaptic cell type and NCD range. This resulted in a convergence value – average number of synaptic inputs to each postsynaptic cell – for each projection Fig. 8*C*. We fixed all connection weights (unitary connection somatic EPSP amplitude) to 0.5 mV, consistent with rat and mouse S1 data (***Hu and Agmon, 2016; Constantinople and Bruno, 2013**).*

To estimate the maximum number of synaptic inputs per region, we made a number of assumptions based on the limited data available (Figs. 8*C* and 1*C*). First, we estimated the average number of synaptic contacts per cell as 8234 by rescaling rat S1 data (***Meyer et al., 2010b**)* based on our own observations for PT cells (***Suter et al., 2013**)* and contrasting with related studies ***(Schüz and Palm, 1989; DeFelipe et al., 2002***); we assumed the same value for all cell types so we could use convergence to approximate long-range input strength. We assumed 80 % of synaptic inputs were excitatory vs. 20 % inhibitory (***DeFelipe et al., 2002; Markram et al., 2015b***); out of the excitatory inputs, 80 % were long-range vs. 20 % local (***Markram et al., 2015b; Stepanyants et al., 2009***); and out of the inhibitory inputs, 30 % were long-range vs. 70 % local (***Stepanyants et al., 2009***). Finally, we estimated the percentage of long-range synaptic inputs arriving from each region based on mouse brain mesoscale connectivity data (***Oh et al., 2014***) and other studies (***Meyer et al., 2010a; Bruno and Sakmann, 2006; Meyer et al., 2010b; Zhang et al., 2016; Bopp et al., 2017***).

Experimental evidence demonstrates the location of synapses along dendritic trees follows very specific patterns of organization that depend on the brain region, cell type and cortical depth ***(Petreanu et al., 2009; Suter and Shepherd, 2015***); these are likely to result in important functional effects (***Kubota et al., 2015; Laudanski et al., 2014; Spruston, 2008***). We employed sCRACM data to estimate the synaptic density along the dendritic arbor – 1D radial axis – for inputs from PO, VL, M2 and OC to layers 2/3, 5A, 5B and 6 IT and CT cell (***Hooks et al., 2013***), and from layer 2/3 IT, VL, S1, S2, cM1 and M2 to PT neurons (***Suter and Shepherd, 2015***) (Fig. 8*D*). To approximate radial synaptic density we divided the sCRACM map amplitudes by the dendritic length at each grid location, and averaged across rows. Once all network connections had been generated, synaptic locations were automatically calculated for each cell based on its morphology and the pre- and postsynaptic cell type-specific radial synaptic density function (Fig. 8*D*). Synaptic inputs from PV to excitatory cells were located perisomatically (50 *μm* around soma); SOM inputs targeted apical dendrites of excitatory neurons (***Naka and Adesnik, 2016; Katzel et al., 2011***); and all inputs to PV and SOM cells targeted apical dendrites. For projections where no synaptic distribution data was available – IT/CT, S1, S2 and cM1 to IT/CT cells – we assumed a uniform dendritic length distribution.

### Model implementation, simulation and analysis

#### Modeling and simulation tools

The model was developed using parallel NEURON (neuron.yale.edu) (***Lytton et al., 2016**)* and NetPyNE (www.netpyne.org) (***Dura-Bernal et al., 2019**),* a Python package to facilitate the development of biological neuronal networks in the NEURON simulator. NetPyNE emphasizes the incorporation of multiscale anatomical and physiological data at varying levels of detail. It converts a set of simple, standardized high-level specifications in a declarative format into a NEURON model. This high-level language enables, for example, defining connectivity as function of NCD, and distributing synapses across neurons based on normalized synaptic density maps. NetPyNE facilitates running parallel simulations by taking care of distributingthe workload and gatheringdata across computing nodes, and automates the submission of batches of simulations for parameter optimization and exploration. It also provides a powerful set of analysis methods so the user can plot spike raster plots, LFP power spectra, information transfer measures, connectivity matrices, or intrinsic time-varying variables (eg. voltage) of any subset of cells. To facilitate data sharing, the package saves and loads the specifications, network, and simulation results using common file formats (Pickle, Matlab, JSON or HDF5), and can convert to and from NeuroML (***Gleeson et al., 2010,2019***) and SONATA ***(Dai et al., 2019)**,* standard data formats for exchanging models in computational neuroscience. Simulations were run on XSEDE supercomputers Comet and Stampede, using the Neuroscience Gateway (NSG) and our own resource allocation, and on Google Cloud supercomputers.

#### Parameter exploration/optimization

NetPyNE facilitates optimization and exploration of network parameters through automated batch simulations. The user specifies the range of parameters and parameter values to explore and the tool automatically submits the jobs in multicore machines (using NEURON’s Bulletin board) or HPCs (using SLURM/Torque). Multiple pre-defined batch simulation setups can be fully customized for different environments. We ran batch simulations using NetPyNE’e automated SLURM job submission on San Diego Supercomputer Center’s (SDSC) Comet supercomputer and on Google Cloud Platform.

#### Local Field Potentials

The NetPyNE tool also includes the ability to simulate local field potentials (LFPs) obtained from extracellular electrodes located at arbitrary 3D locations within the network. The LFP signal at each electrode is obtained using the “line source approximation” (***Parasuram et al., 2016; Buzsáki et al., 2012; Lindén et al., 2013***), which is based on the sum of the membrane current source generated at each cell segment divided by the distance between the segment and the electrode. The calculation assumes that the electric conductivity and permittivity of the extracellular medium are constant everywhere and do not depend on frequency.

#### Firing rates statistics

Firing rate statistics were always calculated starting at least 1 second after the simulation start time to allow the network to reach a steady state. To enable the statistical comparison of the results in Fig. 2 we only included neurons with firing rates above 0 Hz, given that most experimental datasets (***Estebanez et al., 2018; Zagha et al., 2015; Li et al., 2015a**)* already included this constrain. For the statistical comparison in the remaining sections we included neurons with firing rates of 0 Hz, as these were available both in the experimental dataset (***Schiemann et al., 2015**)* and the model. Therefore, the quiet state mean firing rates reported in Fig. 2 (which only included rates > *0Hz*) were higher than those in the remaining sections.

#### Experimental procedures

Details of the experimental procedures used to obtain the data in this study were previously described in (***Schiemann et al., 2015***), including animals and surgery, motion index and motion pattern discrimination, and in vivo electrophysiology and pharmacology. The dataset on cell typespecific in vivo firing rates across states and conditions was collected and previously reported in the same publication. The LFP experimental data reported here was collected during that same study but only a small subset was reported in the experimental paper ((***Schiemann et al., 2015***) Fig. 1).

The experimental LFP data used in Fig. 4 was preprocessed to remove outliers and potential artifacts. The raw LFP data consisted of 30 recordings of varying duration during head-restrained mice locomotion (at different speeds) on a cylindrical treadmill. In order to compare it to the simulated data, the quiet in vivo raw LFP were classified into quiet and movement periods (using the same criteria as in (***Schiemann et al., 2015**))* and then segmented into 4-second samples. We then calculated the LFP power spectral density (PSD) using the Morlet wavelet transform method, normalized within each sample and computed the mean power for five standard frequency bands (delta, theta, alpha, beta and gamma). The resulting dataset of 5-element vectors (normalized power in each frequency band) exhibited high variability: the mean coefficient of variation (CV) across quiet samples was 0.60 and 0.44 for move samples. Therefore we used k-means to cluster the dataset. The quiet condition resulted in one predominant cluster with similar power for all bands (73% of samples), and one with higher gamma power (27% of samples). Conversely, the move condition predominant cluster exhibited significantly higher gamma power (77% of samples), whereas the smaller cluster showed similar power across bands (23%). As expected, the variability within each cluster was significantly reduced compared to the full dataset (large clusters: quiet CV=0.33, move CV=0.32; small clusters: quiet CV=0.31, move CV=0.28). For comparison with the model results we employed the large quiet and move clusters (with over 70% of samples) (Fig. 4). The smaller clusters may correspond to different internal states during behavior, recording from regions/layers with different levels of involvement in the behavior, transition periods, and/or experimental artifacts (e.g. inaccurate segmenting of behavior).

## Acknowledgements

We would like to thank Drs. Zhaga and Estebanez for contributing experimental data for Figure 2. This work was funded by the following grants: NIH U01EB017695 (WWL), NYS SCIRB DOH01-C32250GG-3450000 (SDB,WWL), NIH U24EB028998 (SDB), NSF 1904444-1042C (SDB,WWL), NIH R01 NS061963(GMGS), NIH R01EB022903 (WWL), DARPA N66001-10-C-2008 (WWL), ASAP-020572 (WWL), NIH R01DC012947 (SAN), ARO W911NF-19-1-0402 (SAN) and ARO URAP Supplement. The views and conclusions contained in this document are those of the authors and should not be interpreted as representing the official policies, either expressed or implied, of the U.S. Government or any of its agencies. The U.S. Government is authorized to reproduce and distribute reprints for Government purposes notwithstanding any copyright notation herein. This research was funded in part by Aligning Science Across Parkinson’s [ASAP-020572] through the Michael J. Fox Foundation for Parkinson’s Research (MJFF). For the purpose of open access, the author has applied a CC BY public copyright license to all Author Accepted Manuscripts arising from this submission.

## Acknowledgments

Additional information can be given in the template, such as to not include funder information in the acknowledgments section.

